# Shared and individual tuning curves for social perception

**DOI:** 10.1101/2025.01.19.633772

**Authors:** Rekha S Varrier, Zishan Su, Qi Liang, Tory G Benson, Eshin Jolly, Emily S Finn

## Abstract

Stimuli with light are clearly visual; stimuli with sound are clearly auditory. But what makes a stimulus "social", and how do socialness judgments differ across people? Here, we characterize group-level and individual thresholds for perceiving the presence and nature of a social interaction. We leverage the fact that humans see social interactions—e.g., chasing, playing, fighting—even in very un-lifelike stimuli like animations of geometric shapes. Unlike most prior work that used hand-crafted stimuli, we exploit these animations’ most advantageous property: their visual features are fully parameterizable. Using this property, we construct psychophysics-inspired "social tuning curves" for individuals. We find not only that simple visual features influence social perception, but also that the exact shape of the tuning curve is unique to and stable within each person. Further, individual differences in tuning curves are related to socio-affective traits. Our approach lays the foundation to study how social percepts emerge from interactions between features of a stimulus and features of an individual observer.

## Introduction

A hallmark of the human species is our extraordinary sociality, which depends on reading and responding to others’ behavior in ways that are largely effortless and shared across the population. Yet despite this shared general framework, there are substantial idiosyncrasies in how people perceive, interpret, and react to social information (Abassi & Papeo, 2022; Hull et al., 2023; Nguyen et al., 2019; Ratajska et al., 2020; Varrier & Finn, 2022). Many of these individual differences simply reflect variation in personality traits and social styles. Yet, marked deviations from typical social processing are also central to developmental conditions such as autism (Klin & Jones, 2006; Lisøy et al., 2022; Rasmussen & Jiang, 2019; Vandewouw et al., 2021; Zwickel et al., 2011), as well as mental illnesses such as schizophrenia (Horan et al., 2009; Langdon et al., 2020), paranoia (Castiello et al., 2024) and depression (Kaletsch et al., 2014; Kohler et al., 2011; Krause et al., 2021; Liu et al., 2012).

Social information can take many forms, including linguistic cues, facial cues, and whole-body motion cues. While humans get nuanced information from linguistic and facial cues, motion cues are necessary for some of our most basic evolutionary social behaviors that are conserved across species: e.g., pursuing, evading, playing, fighting, and courting (Barrett et al., 2005; Blythe et al., 1999). Humans are primed to perceive these types of interactions even in very un-lifelike stimuli: for example, when faced with videos of simple geometric shapes moving around the screen, even without prompting, most neurotypical people will construct narratives to explain the shapes’ movements in terms of goals, beliefs, and desires. This highly robust observation dates back at least to Heider and Simmel (Heider & Simmel, 1944) and has since been leveraged to study social perception in a wide variety of contexts and populations (Abell et al., 2000; Gao et al., 2009; Isik et al., 2017; Scholl & Tremoulet, 2000). The effect holds across cultures, suggesting a biological origin (Barrett et al., 2005). Along with related phenomena such as pareidolia (Zhou & Meng, 2020)—the tendency to perceive faces in inanimate objects—this suggests an automaticity to social information processing that belies its typical conceptualization as a high-level cognitive process. Indeed, recent work supports the notion that core components of a social interaction can be extracted by the human visual system using fast, bottom-up processes (McMahon & Isik, 2023), which is likely evolutionarily adaptive for a species that depends heavily on its sociality for survival (Dunbar, 2024; Frith & Frith, 2023).

While using simple geometric-shape animations as experimental stimuli has yielded important insights into behavioral, cognitive, and neural aspects of social perception, most past work using these stimuli has substantial limitations. Studies typically use a small number of manually generated animations handcrafted by human experimenters to be either obviously social or obviously non-social, with no systematic variation or control over visual features (Barch et al., 2013; Castelli et al., 2000; Nguyen et al., 2019). Participants’ responses are then classified as accurate or inaccurate with respect to these “ground truth” experimenter labels.

Furthermore, even given a stimulus deemed “social” by most people, different individuals may be perceiving different *types* of social interactions in that same stimulus; the *nature* of the perceived interaction is rarely probed (and if it is, the experimenters often have a ground truth label in mind —e.g., helping versus hindering) (Hamlin et al., 2007; Rutherford & Kuhlmeier, 2013; Ullman et al., 2009). Together, these practices often produce ceiling effects and compress individual variability in behavior, at least in normative populations, which is unrealistic given that most real-world social scenarios are complex and may engender different interpretations across people.

Here, we exploit a highly advantageous yet hitherto under-used property of such animations—namely, that they are algorithmically controllable and amenable to principles of visual psychophysics—to characterize people’s socio-perceptual tendencies at both the group and individual level. Previous work has taken a psychophysics approach to characterizing percepts of animacy – i.e., whether an entity’s motion appears biological and/or goal-related – by manipulating temporal features (Palmer et al., 2023), change in speed and direction (Tremoulet & Feldman, 2000) and whether they are self-propelled or guided by Newtonian mechanics (Schultz & Bülthoff, 2013). A few previous studies (Gao et al., 2009; Schultz et al., 2005) have also extended this approach to percepts of socialness – i.e., whether two entities appear to be interacting with each other – by manipulating motion contingencies between them. However, this approach remains underexplored as a possible source of individual differences in social perception. We study two processes: (1) how people detect the presence of an interaction and (2) how people discriminate between types of interactions. By parametrically varying motion attributes (Gao et al., 2009), we programmatically generate a large set of animations and use participants’ subjectively reported percepts to construct “social tuning curves” that capture shared trends and individual differences. Throughout, we adopt the perspective that socialness is in the eye of the beholder: in other words, there is no “correct” and “incorrect”; whatever percept is reported is the ground truth for that trial for that participant. We *embrace* ambiguous stimuli—i.e., those that yield high variability in reported percepts—as a feature rather than a bug, as these offer an opportunity to probe the limits of what makes a stimulus social, and how these limits differ for different individuals. We use this framework to show that robust individual differences in socio-perceptual tendencies exist atop group-level trends, that these individual differences are reliable over a period of months, and that they show some relationship to traits indexing real-world social and affective function.

## Methods

### Participants

All data collection and analysis procedures were approved by the Committee for XXX of XXX (withheld for anonymity). All data were collected online (http://www.prolific.com/). We used the following selection criteria: participants had to (1) be fluent in English, (2) have their location set as the USA or UK and (3) not have participated in our previous studies with similar stimuli. For consistency in data quality, all studies – except for the retest sessions where participants (identified using their 24-character Prolific IDs) were invited to complete a second session – were typically launched at 9am Eastern US time on Prolific and closed when the desired sample size was reached (usually about 2pm Eastern US time). The retest sessions were launched 2 months (Mean=71.0 days, SD=5.6 days; detection task) or 1 month (Mean=32.8 days, SD=7.7 days; discrimination task) after their respective first sessions and were left open for about 6 weeks until no participants had completed the task for at least 1 week, with the goal to encourage as many participants as possible to return. Note that the aim of the second sessions was to test the retest reliability (i.e., how similar behavior was on two independent sessions), so the exact time gap between the sessions did not need to be the same, we only required that the session not be so close that we would see effects of perceptual learning or task-specific memory.

#### Stimuli

Stimuli were simple animations live-generated using a custom JavaScript-based software called *psyanim* (https://github.com/thefinnlab/psyanim-2) based on trajectories saved in Google Firebase (https://firebase.google.com/) after quality-checking (see **Supplementary Methods** *Stimuli* è *Quality checks* for more information on the quality checks). Each animation had two circular agents (radius 12 px): one black and one gray, set against a white background in a world of size 800px x 600px (see Fig 1a for a schematic; see recreated recordings of all stimuli here: https://github.com/thefinnlab/psyanim_behav_paper1/tree/master/stimuli/; exact stimuli for the individual experiments will be linked in-text near the description of that behavior). We used circular agents and did not add additional “facingness”-related information (e.g., triangles, eyes) as in some previous studies (Abell et al., 2000; Netanyahu et al., 2021) for two reasons. First, to induce the ambiguity necessary to tease apart individual differences in social perception, we wanted the agents themselves to contain minimal information. Past work (Gao et al., 2009) has shown that predators are evaded more successfully when the agents had such facingness information (i.e., when agents were dart-like compared to discs), suggesting that facingness may enhance percepts of a chase regardless of motion contingencies. Second, while the purpose of facingness details in previous studies may have been to enhance percepts of animacy, other work (Abell et al., 2000) has shown that such details were insufficient and that it was in fact motion patterns that determined whether two triangular agents were seen as social/goal-directed (e.g., chasing, fighting) or non-social and inanimate (e.g., floating in space).

**Fig 1:**
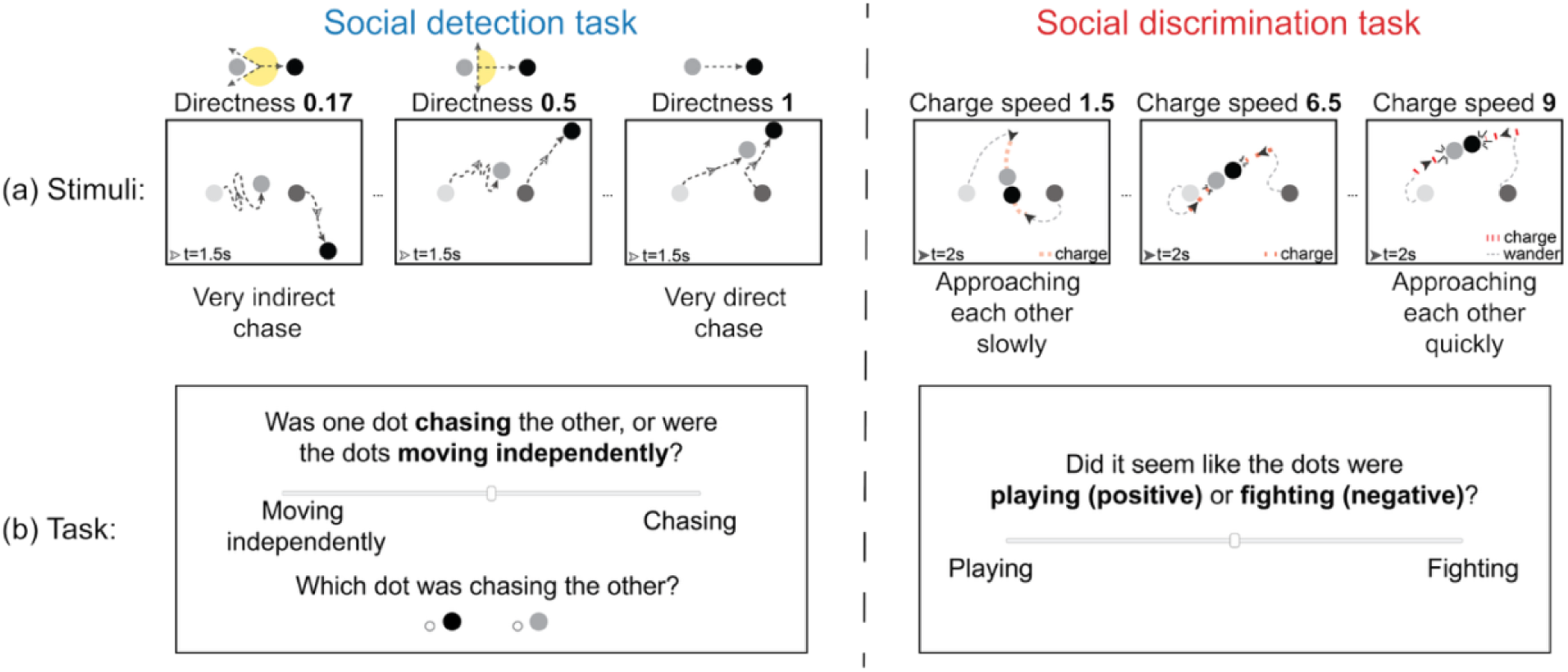
Social detection and discrimination tasks. (a) Static schematics of the animation stimuli for the detection (left) and discrimination (right) tasks to illustrate the effect of varying motion attributes (chase directness and charge speed, respectively) on agents’ trajectories. The two dots at the beginning and the end of each trajectory represent the agents’ positions at the start of the trial and after about 1/4^th^ of the total duration of the animation (at time 1.5s for the detection task and 2s for the discrimination task). For the detection task (left), the yellow zone in the schematic at the top of each directness subplot indicates the range of directions the predator is allowed to take in pursuing the prey (the smaller the range, the higher the directness; directness = (180-subtlety)/180). For the discrimination task (right), the red part of the trajectory indicates when agents switch from wandering to charging at each other. (b) The response screen that was presented following each animation in the main experiments. Both tasks required participants to rate their percept of the preceding animation on a continuous bar, and the detection task (left) additionally asked participants to identify the agent that was doing the chasing. In both experiments, the positions of the slider labels (“Moving independently”/ “Chasing” or “Playing”/ “Fighting”) on the left versus right were counterbalanced across participants.

At the start of an animation, the agents were on either side of the center of the screen (coordinates: 400, 300): left (coordinates: 250, 300) and right (coordinates: 550, 300). In each experiment, the black agent started on the left (gray on the right) in half of the animations, and vice versa in the other half. The animations were 6s (detection task) or 8s (discrimination task) long with a frame rate of 60 Hz. A key difference between this study and most past work on social perception using animations is that we generated our animations purely programmatically using quantifiable, parameterizable motion attributes (Fig 1a). Each experiment consisted of 7 levels of stimuli where one motion attribute of interest varied linearly across levels while all other attributes were held constant. These attributes were chosen such that people’s perception of a social scene on a scale from the most non-social to most social (detection task) or from most playful to most aggressive (discrimination task) varied along the attribute of interest. In the following sub-sections, we describe these motion attributes as well as the animations in more detail.

#### Detection task

To manipulate percepts as to the presence versus absence of a social interaction (detection task), we relied on the motion attribute *chase directness*, which governs the fidelity with which one agent (the “predator”) chases the other agent (the “prey”). This attribute was originally described by Gao et al. (Gao et al., 2009), where it was called “chase subtlety” and was found to robustly influence people’s ability to detect social interactions (in particular, chases). Chase subtlety was originally defined in angles, i.e., by how many degrees a predator could deviate from a perfect heat-seeking path between itself and the prey at each time step. Thus, a *chase subtlety* of 0° would indicate a very direct chase, a *chase subtlety* of 90° would indicate a somewhat noisy chase where the predator can go off-path by up to 90° clockwise or 90° counterclockwise, and *chase subtleties* > 90° would indicate very noisy chasing behaviors where the predator can occasionally even move away from the prey. Here, we reversed and normalized the subtlety angle to derive *chase directness* 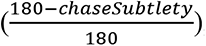, such that the higher the directness, the more obvious (detectible) the chase. Details on how this was implemented in *psyanim* are below.

### Chase animations

In the animations we generated, one agent is the predator and the other is the prey. Predator/prey assignment was counterbalanced across stimuli in terms of both start position (left/right of the center) and color (gray/black). The predator was programmed to chase the prey at varying levels of *chase directness* and the prey was programmed to flee from the predator when it was within a certain radius. When the predator agent was beyond its field of view (or the distance between them was greater than the “*safety distance*”), the prey agent simply wandered around the screen. We included the wandering behavior to prevent the fleeing behavior from looking too obvious, so that people did not make decisions purely based on the prey. All variable attributes other than *chase directness* governing the motions of the predator and prey were held constant over all animations (i.e., over all levels of *chase directness*). The stimuli used for session 1 chase detection experiments are <u>here</u> (https://github.com/thefinnlab/psyanim_behav_paper1/tree/master/stimuli/detection_subtlety/Chase), and those for session 2 experiments are <u>here</u> (https://github.com/thefinnlab/psyanim_behav_paper1/tree/master/stimuli/detection_subtlety/Chase_set2). The relevant attributes are described in the **Supplementary Methods** section *Stimuli* (sub-sections *Chase animations* and *Wander behavior*).

### “Invisible chase” control condition

With this control, we sought to rule out an alternative possibility for how *chase directness* might influence socialness perception that is less related to a chase *per se* and more related to general motion contingency between the two agents. Specifically, observers may notice that the predator and prey trajectories are linked more tightly in time at higher *chase directness* (where immediately after the prey changes direction, the predator too will change direction) than at lower *chase directness* (where the predator will not change direction as quickly and obviously upon the prey changing direction). Participants may simply be using this heuristic—i.e., whether the prey changing direction prompts the predator to change direction—instead of the actual chase between the predator and prey. To test for this possibility, inspired by Gao et al. (Gao et al., 2009), in a subset of behavioral experiments we included an additional set of control stimuli, where the actual prey was initialized at a randomly chosen location on the screen for each animation but was made invisible. The predator and a visible “mimicking” agent each started at one of the two regular starting locations (left and right of center as described at the start of the **Stimuli** section). The mimicking agent copies the true prey’s trajectory but with a 180° rotation (i.e., if the invisible prey moves up and to the right, the mimicking agent will move down and to the left). The animations used for this study are <u>here</u> (https://github.com/thefinnlab/psyanim_behav_paper1/tree/master/stimuli/detection_subtlety/Mimic_cheat). For illustrative purposes, the invisible (true) prey is shown in yellow in these exemplars (participants never saw the yellow dots!). See **Supplementary Methods** section *Stimuli* è *Invisible “chase” control* for the stimulus attributes of the mimicking agent in psyanim.

#### Discrimination task

To study how people discriminate between positive and negative social interactions, we varied the attribute *charge speed* in a novel social interaction scene that is different from the chase detection task discussed above. This scene was inspired by the presence of physical contact in the real world for common positive as well as negative interactions (e.g., hugs, high-fives, physical fights) (Neri et al., 2006) and how speed can be a clue to valence, with slower movements being usually perceived as more peaceful/positive, and faster movements as more aggressive/negative (Blythe et al., 1999).

When generating these animations, both agents were set to have the same goal: to wander for a certain period and then charge at the other agent at the predetermined *charge speed* which varies between 1.5px/frame and 9px/frame (stimuli in Fig 1a and <u>here</u> (https://github.com/thefinnlab/psyanim_behav_paper1/tree/master/stimuli/discrimination_playfight). Once one agent initiates a charge, the other agent will respond after a short delay; following contact, both agents will return to wandering. For more details on the stimulus attributes, see **Supplementary Methods** (*Stimuli è Discrimination task*).

#### Animacy cover story

Attributing intentions to moving shapes entails judgments of two related yet distinct features – animacy and socialness. Animacy is a widely used concept that applies to entities that are considered alive (Trompenaars et al., 2021). These entities exhibit signs of self-propelled, non-Newtonian motion by seeming to engage in goal-directed behavior (Schultz & Frith, 2022) and responding to their surroundings (e.g., changing speed or direction to avoid an obstacle). In our view, animacy is necessary but not sufficient for socialness: animacy can be detected in displays of single agents, in which by definition there is no social interaction present, but in multi-agent displays, to the extent that agents are perceived to be socially interacting, they must also be perceived as animate (i.e., as possessing a mind that would be motivated to engage in social behavior). Our goal here was to isolate the concept of *socialness* above and beyond animacy.

Because differences in percepts of animacy might confound judgments of socialness, to encourage uniform perception of animacy across both animations and participants to the extent possible, we provided a cover story that the agents (dots) represented children in a public park. A cover story is a more effective way to ensure a uniform level of animacy across all stimuli amongst all participants in comparison to physical features such as adding facingness information to agents (e.g., triangular agents) or eyes – past work has shown that the anthropomorphism in the Heider and Simmel stimuli was preserved even when the shapes were distorted (Berry et al., 1992) and that the perception of the same agents as animate or inanimate depends on motion patterns, not features of the agents themselves (Abell et al., 2000). We chose this cover story specifically because it was believable (people care about children’s privacy), relatable (children in playgrounds are a common sight) and helps us to test individual differences in various behaviors (chasing, moving independently, playing, fighting) with the same cover story without changing people’s interpretations of what the entities could have represented.

However, as we discuss later, this did introduce a bias towards perceiving playful interactions in the discrimination task. This was the exact story participants received: “*We recently videotaped a public park where nearby children go, with the goal of capturing the essence of children’s behaviors within a familiar park setting. To protect the identities of these young people, we used an algorithm that represents **a pair of children as two dots**, each tracing the path of an individual child.”* We used this cover story to set the context in all experiments except in the early pilot experiments, since the goal of the latter was to evaluate how participants spontaneously interpret the animations without any prompting or context.

### Data collection

All pilot and main experiments were programmed and run using the *jsPsych* platform (Leeuw et al., 2023) with a custom plugin (https://github.com/thefinnlab/psyanim-2) to present the *psyanim* animations.

### Experimental design

We first conducted a set of pilot experiments to verify that our algorithmically generated animations can in fact spontaneously evoke percepts that fall approximately along the intended axes from non-social to social (detection task) or playful to aggressive (discrimination task).

Participants watched the animations and made free text responses of 1-2 sentences. Details of the pilot experiments are in the **Supplementary Methods** (section *Pilot experiments (open-ended responses)*). These experiments showed that even without explicit constrained rating scales, our stimuli could spontaneously evoke percepts along the intended axes, which in turn gave us the confidence to move forward with our main experiments that did use explicit, constrained rating scales to quantify perception more precisely.

### Main experiments

#### Main experiment design

Our first two main experiments consisted of only detection or only discrimination trials, respectively, while the third main experiment was a mixed-task design in which the same participants performed both the detection and discrimination tasks. We used the same cover story described above (about the dots being children in a public park) in all the main experiments.

For the detection study, there were 84 trials per participant (12 trials per *chase directness* level x 7 levels) with 7 optional breaks (one every 12 trials). In the control experiment that also included the invisible chase control condition, there were 6 stimuli at each motion attribute level (2 conditions x 7 stimulus levels x 6 trials per level). After each trial, participants gave two responses: (1) they rated, on a continuous scale, to what degree one of the two dots was chasing the other versus moving independently (with the location of the labels on the left versus right extremes of the scale kept constant within participant, but counterbalanced across participants), (2) they identified the dot that was chasing the other, in a two-alternative forced choice. Both questions were presented on the same page, and the page timed out in 10s. The instruction specific to this task (after the cover story was presented) was as follows: “*Some videos depict a situation in which **one dot is chasing the other**; other videos depict a situation in which the dots are **moving independently**. You will be asked to **rate how much you think the dots are interacting (meaning one dot is chasing the other) versus moving independently**. For all videos, you will also be asked to determine **which dot was chasing the other**. If there was no chase in a particular video, just make a guess.*”. Participants performed one practice trial before the experiment began.

For the discrimination study, there were 70 trials per participant (10 trials per *charge speed* level x 7 levels) with 7 self-timed breaks (one every 10 trials). After each trial, participants rated on a continuous scale to what degree the dots were engaged in a positive (playing) versus negative (fighting) interaction (as with the detection task, the label positions were counterbalanced across participants). This response page also timed out in 10s. The instruction specific to this task (after the cover story was presented) was as follows: “*Some videos depict a situation in which the children are engaged in a **positive interaction (e.g., playing)**; other videos depict a situation in which the children are engaged in a **negative interaction (e.g., fighting)***.

*After each video, you will be asked to **rate to what extent the children (dots) were engaged in a positive or negative interaction** on a continuous bar. There are no right or wrong answers here, so if you are unsure, just guess!”*. Participants performed two practice trials before the experiment.

In the third main experiment (mixed-task design study), detection and discrimination animations were presented in six interleaved blocks: block sequence with 14 trials each (121212 or 212121, where 1= detection task and 2 = discrimination task; one of the two block sequences was randomly selected for each participant. Here too, there were 84 trials in total: 42 detection trials (6 at each level of *chase directness*) and 42 discrimination trials (6 at each level of *charge speed*). The 42 animations from each task (detection or discrimination) were randomized *across* the 3 blocks of the task. Participants performed two practice trials each for the detection and discrimination experiments at the start of the experiment.

The primary task portion of all three experiments (detection, discrimination, or mixed-task design) lasted approximately 15-20 min. At the end of the primary task portion, we presented participants with the trait questionnaires described below under **Trait measures**. The sequence of the questionnaires was counterbalanced across participants.

#### Main experiment analysis

Data analysis was similar across all main experiments unless specified otherwise.

#### Separate detection and discrimination experiments

##### Group-level analyses

We first analyzed data at the group level to ascertain shared tendencies in how motion attributes affect social percepts. Participants’ ratings were coded on a 0–1 scale: (i) detection task: *moving independently* = 0, *chasing* = 1; (ii) discrimination task: *playing* = 0 and *fighting* = 1. We used linear mixed-effects analyses to quantify the effect of each motion attribute while controlling for confounding variables using the following model: 𝑟𝑎𝑡𝑖𝑛𝑔 ∼ 𝑚𝑜𝑡𝑖𝑜𝑛_𝑎𝑡𝑡𝑟𝑖𝑏𝑢𝑡𝑒_𝑙𝑒𝑣𝑒𝑙 + 𝑚𝑒𝑎𝑛_𝑑𝑖𝑠𝑡 + 𝑡𝑟𝑖𝑎𝑙_𝑛𝑢𝑚𝑏𝑒𝑟 + (1|𝑠𝑢𝑏_𝑖𝑑) + (1|𝑠𝑡𝑖𝑚_𝑖𝑑). Here 𝑟𝑎𝑡𝑖𝑛𝑔 refers to participant responses indicating the level of socialness (degree of *chasing*) or aggressiveness (degree of *fighting*). The term 𝑚𝑜𝑡𝑖𝑜𝑛_𝑎𝑡𝑡𝑟𝑖𝑏𝑢𝑡𝑒_𝑙𝑒𝑣𝑒𝑙 refers to degree of either *chase directness* (detection) or *charge speed* (discrimination) and could take one of 7 levels. Additional terms are (i) 𝑚𝑒𝑎𝑛_𝑑𝑖𝑠𝑡, the distance between the two agents averaged across all frames; past work has shown that agents that are closer together are more likely to be perceived as interacting (Rasmussen & Jiang, 2019); (ii) trial number, indicating serial order over the course of the experiment (to check for any drift in ratings over time); and random-effects terms for (iii) participant identity (𝑠𝑢𝑏_𝑖𝑑) and (iv) specific animation identity (𝑠𝑡𝑖𝑚_𝑖𝑑). For the predator-identification question in the detection task, we ran a logistic regression model with accuracy (0/1) as the dependent variable and the same main- and random-effects predictor terms as above.

### Individual-level analyses

In addition to showing group-level effects of these motion attributes on social perception, we also expected to see individual differences in how social perception varies *across* the motion attribute levels – for instance, some participants’ perception may change linearly with the motion attribute levels whereas others’ perception may follow a non-linear sigmoid pattern. Similarly, some participants may perceive social or aggressive stimuli at lower *chase directness* or *charge speed* levels than others. Hence, we next analyzed data at the individual level to determine the extent to which participants differed in their socio-perceptual tendencies, how stable these differences were across sessions, and how detection and discrimination tendencies covary with one another and with other socio-affective traits. Our primary approach to analyzing individual-level data was to compute single-subject “tuning curves” for detection and/or discrimination behavior. We averaged each participant’s responses at each motion attribute level (*chase directness* or c*harge speed* level for detection and discrimination tasks, respectively). For each participant, we plotted motion attribute level (normalized to a 0-1 range; x-axis) against average rating across animations at that level (y-axis). Similar to the group-level results shown in Fig 3, visual inspection suggested that individual detection ratings followed a sigmoid shape, while discrimination ratings followed a more linear trend. We empirically tested both sigmoid and linear fits to evaluate which one better fit each type of data and verified this pattern: the Akaike Information Criterion (AIC) – which indicates which of two models fits the data better (lower AIC indicates better fits) – was lower for the sigmoid fits in the detection task (mean difference sigmoid – linear fits ≤ –9.4, *p* < .001 based on paired *t*-test in sessions 1 and 2 of the detection experiments; AIC_sigmoid_ – AIC_linear_ < –5 in 90% and 86.8% of all participants in session 1 and session 2, respectively) and lower for the linear fits in the discrimination task (mean difference sigmoid – linear fits ≥ 7.21, *p* < .001 in sessions 1 and 2 of the discrimination experiments; AIC_linear_ – AIC_sigmoid_ < –5 in 90.7% of all participants in both session 1 and 2; supplementary Fig S2). Hence, we used sigmoid fits to characterize each participant’s response data in the detection task and linear fits to characterize responses in the discrimination task.

The sigmoid curve-fitting equation was : 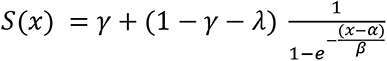, where 𝑥 = the value of the motion attribute (*chase directness* or *charge speed*); 𝛾 and 𝜆 = the lower/upper asymptote of the curve; 𝛼, 𝛽 = center, slope. The linear equation was: 𝐿(𝑥) = 𝑐 + 𝑚 ∗ 𝑥 , where 𝑥 = the motion attribute (*chase directness* or *charge speed*); *c* = the lower intercept of the line; *m* = slope. For both functions, we calculated several key parameters from the fitted curve of each participant:

***Shifts from extremes***. Lower bias 𝑙𝑏 and upper bias 𝑢𝑏 (lower and upper intercepts, respectively) reflect ratings at the lowest and highest levels of the motion attribute — in other words, how close to the extremes of the rating scale a participant is willing to go. For linear fits, 𝑙𝑏 = intercept *c*.

***Bias***. This parameter was derived from the above-mentioned bias terms (𝑙𝑏 and 𝑢𝑏) as a comprehensive summary of people’s bias that also factors in apparent biases due to differences in overall confidence or perceptual vividness. The *bias* term thus measures to what degree people avoid the lowest end of the scale relative to how much they avoid the two ends of the scale in general (which could reflect lower confidence overall or a less intense effect of stimuli overall).

We quantified this as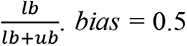 means that there is no bias towards one end of the scale, *bias* < 0.5 means that people are more biased towards the lower end, and *bias* > 0.5 means that people are more biased towards the upper end. In the detection task, *bias* > 0.5 can be interpreted as a predisposition to see things as social (“chasing”) more than non-social (“moving independently”); in the discrimination task, *bias* > 0.5 can be interpreted as a predisposition toward seeing things as more like “fighting” than “playing”.

***Midpoints***. Two types of midpoints can be derived from each curve: an objective midpoint (𝑥_𝑜𝑏𝑗_ = 𝑓^−1^(𝑆(𝑥) = 0.5)) and a subjective midpoint (*subj_center*, 𝛼). The objective midpoint is the motion attribute value at which the participant’s rating crosses the absolute midpoint of the rating scale (0.5 for all participants). The subjective midpoint is the motion attribute value at which the participant’s rating crosses the halfway point of *their own* behavioral curve (e.g., for a participant whose ratings vary between from 0.4 and 0.8, their mid-point would be the motion attribute level at which the tuning curve crosses 0.6). Here we mostly focus on the objective midpoint, which we call the point of subjective equivalence (*PSE*) because it is similar in spirit to *PSE* as defined in traditional visual psychophysics work (i.e., the stimulus feature level at which options A and B are equally likely in a two-alternative forced-choice discrimination task (Gescheider, 1985)).

***Range***. The distance between the lowest and highest ratings on the y-axis. This parameter can be interpreted in multiple ways: it quantifies how much of the total response scale participants use, how much participants differentiate between stimuli at extreme (lowest and highest end) motion attribute levels, and except in cases of strong response biases, might reflect how confident people are in their percepts (especially at the lowest and highest motion attribute values). This is defined as 1 − 𝑙𝑏 − 𝑢𝑏.

***Sigma (σ; sigmoid fit only)***. Sigma or the inverse of the slope 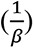 determines the steepness of the sigmoid curve during its transition from perceiving something as “moving independently” (at lower *chase directness* levels) to “chasing” (at higher *chase directness* levels) in the detection task. A smaller sigma (or a higher slope) indicates that participants perceive something as more social (Δ_rating_) with the smallest change in the sensory evidence (change in *chase directness* or Δ_chase_directness_) ⎯ this can be interpreted as people exhibiting higher confidence when rating the intermediate, ambiguous stimuli. A higher sigma (or lower slope) indicates that participants need a lot more sensory evidence (Δ_chase_directness_) to rate something as more social (Δ_rating_) ⎯ this may reflect lower confidence and/or a more gradual, evidence-based shift in the perceptual intensity of stimuli at the ambiguous middle levels.

We plotted covariance matrices (Pearson *r*) between the various parameters within the detection and discrimination tasks. Based on these covariances as well as test-retest reliability of the curve parameters (described in detail below), in what follows, we focus on three main curve parameters that are both relatively reliable and not highly collinear with one another: *PSE*, *range* and *bias*. These terms are illustrated in Fig 2. Since *range* covaries tightly with *PSE* in the linear fits specifically and it is not possible to change *range* without changing the *PSE*, we only use *PSE* and *bias* for the discrimination task. Approximations were made to *PSE* ∉ [0,1]. In participants with a strong social bias (detection task) or aggressiveness bias (discrimination task) at the lowest motion attribute level (*lb* > .5, *ub* < .5; see Fig 2), *PSE* < 0, = ∞ or it is not computed; hence, this was approximated to 0 (detection task: N = 11 out of 312 total in session 1, N = 10 out of 240 total in session 2,; discrimination task: N = 1 out of 319 total in session 1, N = 1 out of 269 total in session 2, . In participants with a strong non-social bias (detection task) or a playfulness bias (discrimination task) at the highest motion attribute level (*lb* < .5, *ub* >.5), *PSE* > 1, = ∞ or it is not computed; this too was approximated to 1 (detection task: N = 0 for detection task session 1 & 2; discrimination task: N = 1 out of 319 total in session 1, N = 0 in session 2).

**Fig 2:**
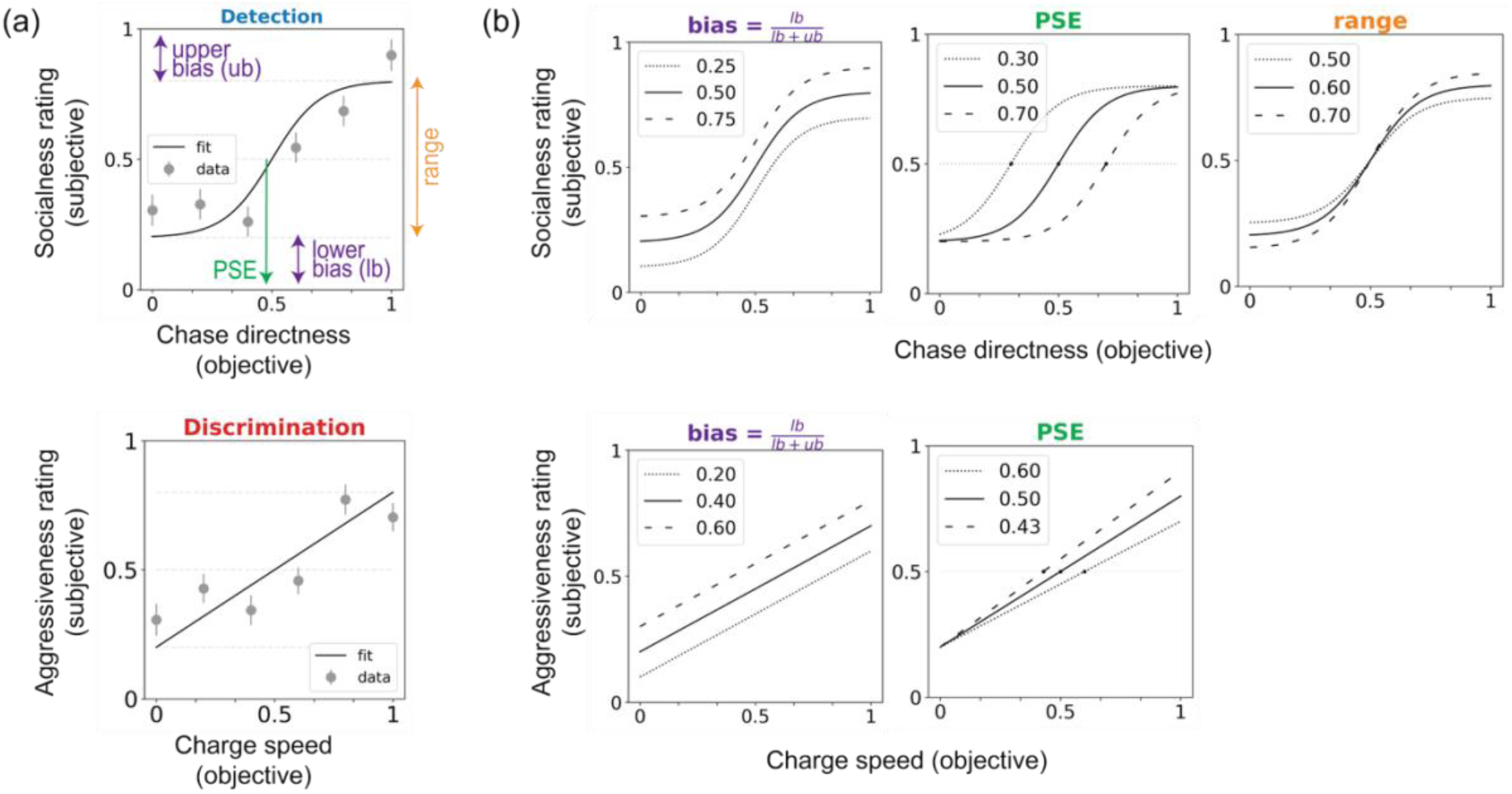
Fitting social tuning curves to individuals’ reported percepts. (a) We fit sigmoid (detection experiments; top) or linear (discrimination experiments; bottom) curves to individual-participant rating data and used the resulting curve parameters to characterize individual participants. (b) Schematics of how each parameter can vary across participants. The upper rows show sigmoid fits as used for the detection task, and the lower rows show the linear fits used for the discrimination task. Each plot shows curves for three different hypothetical participants.

**Fig 3:**
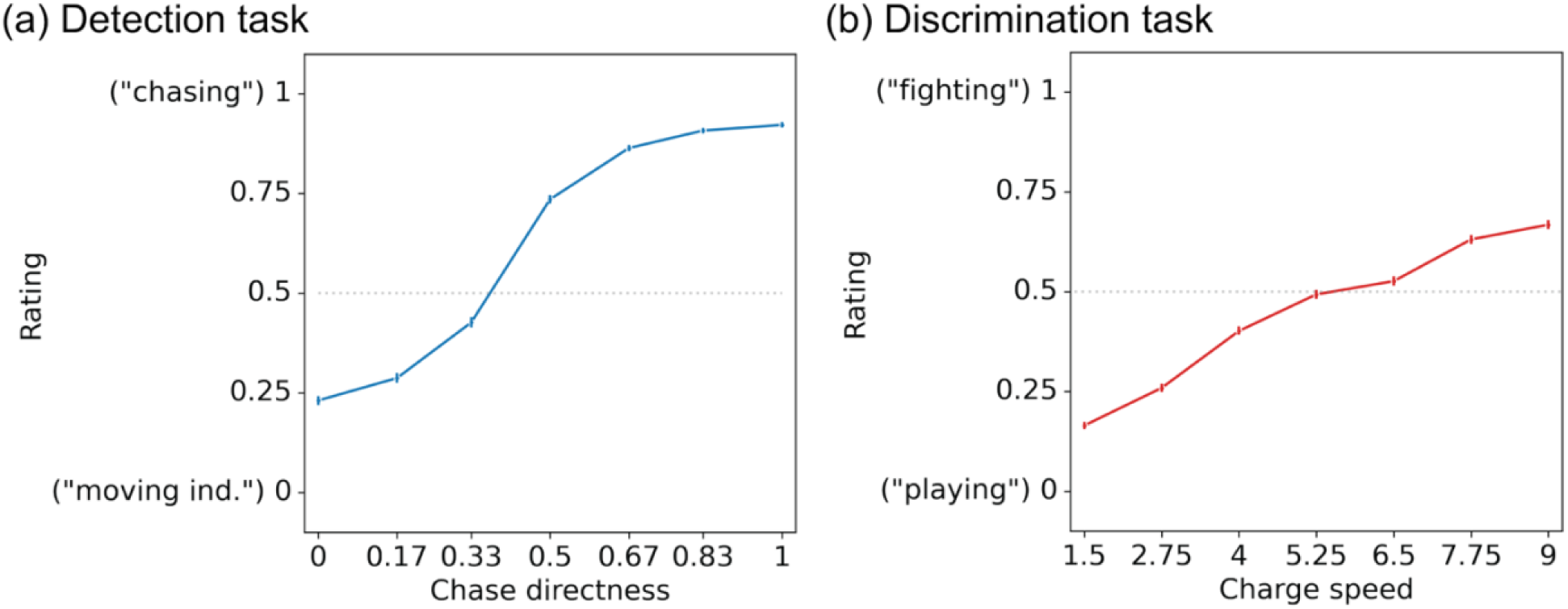
Group-level behavior on the detection and discrimination tasks. (a) As chases become more direct, observers were more likely to report percepts closer to the “chasing” (social interaction) end of the scale. At less direct chases, ratings were lower (i.e., closer to “moving independently [“moving ind.”]). (b) As the charge speed (speed at which agents charge at each other) increased, interactions were perceived to be more aggressive (closer to the “fighting” end of the scale). At lower charge speeds, ratings were closer to “playing”. N = 312 and 319 in panels (a) and (b), respectively. Error bars represent the 95% confidence interval.

Lastly, *range* < 0 suggests a flipped curve (e.g., socialness rating is highest in purely wandering stimuli and lowest at *chase directness* = 1) – this was rare and hard to interpret, hence these participants too were excluded from further analyses (detection task: N = 0 for detection task session 1 and 2; discrimination task : N = 8 out of 319 total in session 1, N = 5 out 269 total in session 2).

We tested the robustness of each person’s tuning curve by using parameters calculated on data from their first session to fit the same participant’s data in their second session 1–2 months later (and vice versa) and calculating the residual normalized root-mean squared error (NRMSE) between the curve and the true data points. We used a paired *t*-test to compare the NRMSE from this within-participant fit to the NRMSE from the mean across-participant fit (calculated by averaging the curve parameter values across all other participants in the same session. This quantified the extent to which an individual’s tuning curve offered a better prediction of their own held-out data than generic tuning curves based on group data. Further, we calculated the intra-class correlation coefficient (ICC) of each curve parameter of interest to measure test-retest reliability across the two sessions (ICC2, single random raters, as implemented by the Python package *pingouin*).

#### Mixed-task experiments

In these experiments, the same participants performed both the detection and discrimination tasks. Data extraction and curve-fitting was performed similarly to the separate detection and discrimination experiments described above; curve-fit patterns were overall similar to the separate detection and discrimination experiments – a sigmoid function fit detection task data better (mean difference sigmoid – linear fits = –7.84, *p* < .001 based on paired *t*-tests, AIC_sigmoid_ – AIC_linear_ < –5 in 82.3% of participants) and a linear function fit discrimination task data better (mean difference linear – sigmoid fits = –3.44, *p* < .001 based on paired *t*-tests, AIC_linear_ – AIC_sigmoid_ < –5 in 77.2% of participants). For each participant, we obtained tuning curves for both detection and discrimination. Similar to the separate experiments, *PSE* for people with high non-social (playfulness) or social (aggressiveness) biases was set to 0 (detection: N = 9, discrimination: N = 2, out of 279 total) or 1 (detection: N = 0 , discrimination: N = 1, out of 279 total) and participants with range < 0 were excluded (detection: N = 0, discrimination: N = 3, out of 279 total; details in the previous sub-section). We studied how socio-perceptual tendencies in the two tasks relate by calculating the Pearson correlation coefficient between all possible pairs of curve parameters across the detection and discrimination tasks. Here, we restricted our analyses to only the most robust curve parameters: *bias*, *range* and *PSE* for the detection task, and *bias* and *PSE* for the discrimination task (5 total).

Traits were scored as described under **Trait measures** below, giving us 14 dimensions in total (5 for AQ, 2 for PANAS, 5 for NEO-FFI, 1 for loneliness and 1 for number of friends). To study how traits relate to behavior on our task, we performed multiple linear regressions fitting the curve parameters to the trait dimensions. We first combined two terms that measure similar properties — *PSE* and *bias* — in a composite index in both the detection and discrimination tasks (separately). We created this composite index to reduce redundancy, since these terms were moderately (*r* = −0.44) and strongly (*r* = −0.70) correlated in the detection and discrimination tasks, respectively (Fig 5), and because we did not have strong hypotheses about how they might differentially relate to traits. In the detection task, this index reflects sensitivity to social information in that both higher *bias* and lower *PSE* reflect the tendency to report social interactions at lower social evidence levels. In the discrimination task, this index reflects people’s tendency to report interactions as aggressive, in that higher *bias* means higher aggressiveness ratings even at the lowest *charge speed* and lower *PSE* means lower *charge speed* at which animations are first perceived as aggressive. Since *bias* is measured on the y-axis (ratings) and *PSE* is measured on the x-axis (stimulus level), both measures were first normalized across participants by z-scoring; *PSE_z* was then subtracted from *bias_z*. This term will be henceforth referred to as *bias – PSE*. Next, we performed 4 multiple linear regressions: the 14 trait dimensions were fitted to the *bias – PSE* and *range* parameters from the detection and discrimination tasks. To assess statistical significance of each predictor (trait), for each of the 4 models, we bootstrapped (sampled with replacement) participants 1000 times and calculated the 95% confidence interval (*CI*) for each of the 14 regression coefficients. Traits whose 95% *CI*s do not cross 0 are deemed significant predictors of the respective curve parameter. To further assess overall goodness of each model, we performed permutation tests by scrambling the curve parameters *(bias – PSE* and *range*) with respect to the trait scores across participants 10,000 times and computed the fraction of permuted models whose *R^2^* value was higher than that of the true model (*p*-value).

### Trait measures

We chose individual-difference measures of interest based on past findings that behavior on social perception and cognition tasks often differs between populations (e.g., neurotypical versus autistic, people with depression versus without) and/or covaries in the normative population with socio-affective and personality traits. Past work has shown that people scoring higher on autism-like phenotypes are less likely to detect intentions and interactions in social animation displays (Klin & Jones, 2006; Lisøy et al., 2022; Rasmussen & Jiang, 2019; Vandewouw et al., 2021; Zwickel et al., 2011), while people with more internalizing symptoms (related to anxiety and social withdrawal) and a higher desire for social connection are *more* likely to detect intentions and interactions (Epley et al., 2008; Gardner et al., 2005; Powers et al., 2014; Tomova et al., 2020; Varrier & Finn, 2022). Other studies have associated depression with impaired emotion recognition of social stimuli (hypersensitivity to negative cues, hyposensitivity to positive cues) (Kaletsch et al., 2014; Kohler et al., 2011; Krause et al., 2021; Liu et al., 2012). We assessed autism-like traits with the autism quotient (AQ) questionnaire (Baron-Cohen et al., 2001), loneliness with the UCLA loneliness scale (Russell et al., 1978), and general affect with the Positive and Negative Affect Schedule (PANAS) (Watson et al., 1988). We also administered the NEO five-factor inventory for multidimensional personality (NEO-FFI; a measure of the “Big Five”) (Franić et al., 2014). Finally, we asked participants to state the number of close friends they had, since this metric provides additional information about participants’ real-world social tendencies.

Thus, our final battery thus consisted of five entities: (i) AQ, (ii) PANAS, (iii) NEO-FFI, (iv) UCLA loneliness scale and (v) the self-reported number of friends. Details of the questionnaires are given in the **Supplementary Methods** section *Trait measures*.

Correlations among the 14 trait dimensions are shown in supplementary Fig S7.

### Code and data availability

All stimuli, data and code will be available upon publication on GitHub.

## Results

We systematically studied how people perceive the presence and nature of a social interaction using sets of algorithmically generated animations. Each animation consisted of two agents—one gray and one black circle—that were programmed to move in certain ways with respect to one another. Critically, the animations varied parametrically along one motion attribute and were controlled for all other low-level visual features. While our intentionally simple visual displays may not evoke social percepts as spontaneously as other hand-crafted animations used in past work (e.g, Frith-Happé animations; Castelli et al., 2000), this is by design, since a high degree of ambiguity with respect to the property of “socialness” is necessary to bring out individual differences in social perception. We conducted three sets of experiments: (1) detection studies, in which participants rated the extent to which the agents appeared to be interacting (i.e., one agent chasing another) versus moving independently, (2) discrimination studies, in which participants rated the extent to which the agents appeared to be fighting versus playing, and (3) a mixed-task study where participants performed both the detection and discrimination experiments (see stimuli <u>here</u> and details in Fig 1). Below, we describe group-level and individual behavioral patterns for both social detection and discrimination, as well as how these two behaviors compare to one other and to self-reported social and affective traits. The group-level analyses are a replication of past work (detection task) or based on intuitions about speed (discrimination task). After validating group-level effects, we study individual differences, namely robustness of response patterns over multiple sessions and how the patterns relate to behavioral traits.

### Simple motion attributes influence percepts of both the presence and nature of social interactions

Our first goal was to validate that the motion attributes of interest, *chase directness* (which determines the fidelity with which one agent – the predator – moves toward the other agent – the prey) and *charge speed* (which determines how fast the agents approach one another) influence social perception in the same general way at the group-level. Specifically, we tested whether in the detection task, we could replicate previous results by Gao et al. (Gao et al., 2009) that more direct chases will be perceived as more obviously social and less direct chases will be perceived as agents moving independently. In the discrimination task, we asked whether in line with our intuitions based on past work (Blythe et al., 1999), agents charging at each other slower will look more playful and those charging at each other faster will look more aggressive/fight-like.

### Pilot experiments

We first performed pilot experiments to test that these animations could spontaneously evoke percepts along the intended continua without any explicit prompting. In these experiments, participants watched animations and gave free-response text descriptions, which we quantified using tools from natural language processing. Indeed, we found that varying *chase directness* elicited percepts along a continuum from moving independently (non-social) to chasing (social), while varying *charge speed* elicited percepts along a continuum from positive/playing/low arousal to negative/fighting/high arousal (see **Supplementary Results** and supplementary Fig S1).These results showed that our parametrized stimuli generated via *psyanim2* could indeed evoke varying levels of socialness, which gave us confidence to use these stimuli for our main experiments, in which we replaced free responses with these continua as predetermined rating scales.

### Detection experiments

In our main social detection experiments (see Table 1 for sample sizes and other relevant information), after watching each animation, participants (1) rated how social it was on a continuous scale ranging from “moving independently” to “chasing” (henceforth referred to as “socialness rating”), and (2) identified the predator agent by color. In line with our expectations and past work (Gao et al., 2009), we found that as *chase directness* increased, ratings shifted towards “chasing” (*b* = 0.805, *p* < .001; Fig 3a). This shift follows a non-linear (sigmoid) pattern overall, as evidenced by the lower AIC in all the detection experiments for sigmoid compared to linear fits (supplementary 2a; mean difference in model fits AIC_sigmoid_ – AIC_linear_ ≤ –7.84; *p* < .001 in the paired *t*-tests in all the detection experiments). Social perception did not seem to change with time (indexed as the trial number; *b* = –0.001, *p* = .779), suggesting that there was no measurable “drift” in ratings toward more social or more non-social over the course of the experiment. We also observed that, similar to subjective ratings, predator identification accuracy increased as chases became more direct (logistic regression *b* = 4.735, *p* < .001; supplementary Fig S3a); this result further confirmed that participants were, on average, experiencing the chase in the expected way based on the generating algorithm (i.e., they correctly perceived its directionality, especially in the case of the more direct chases).

**Table 1:**
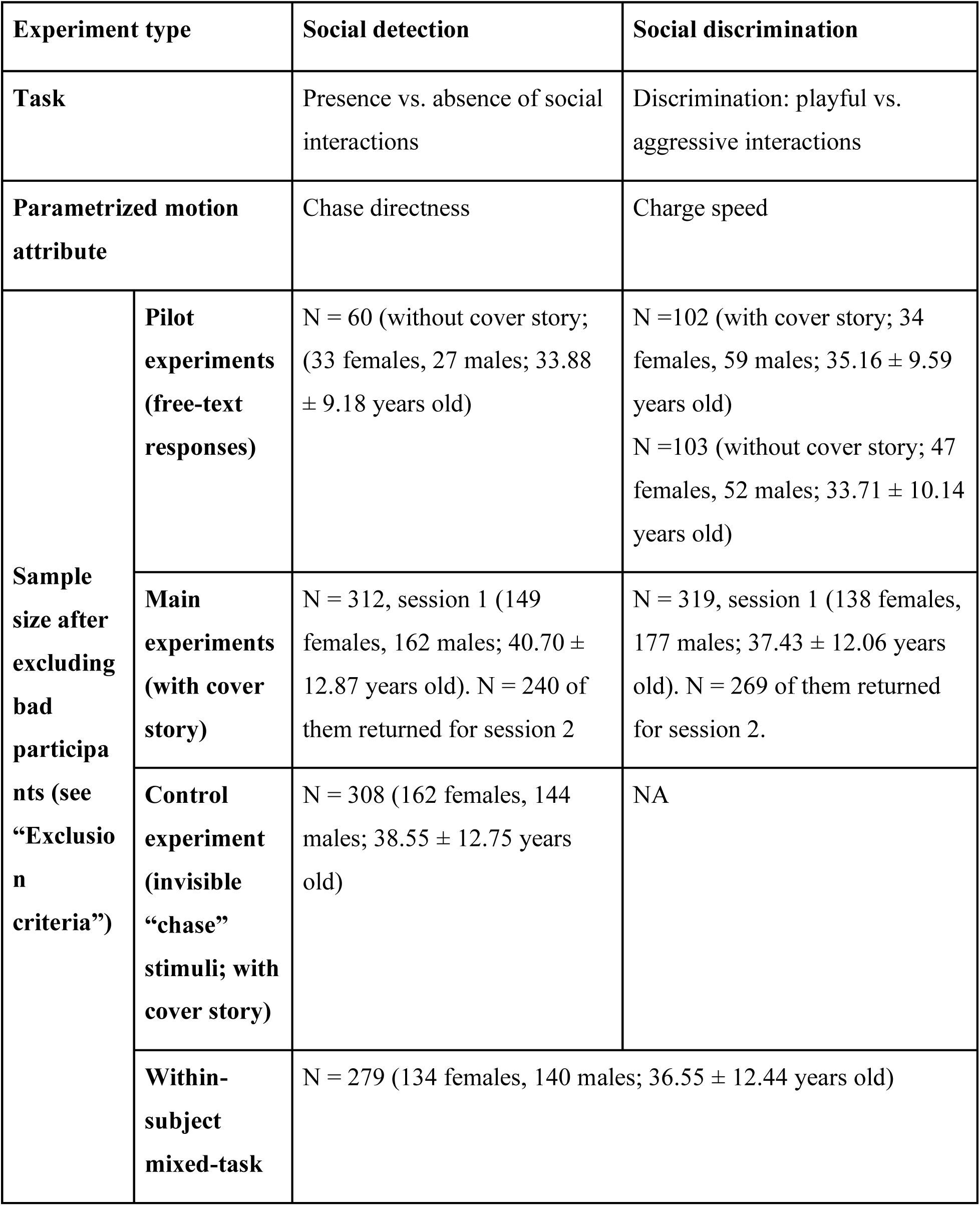
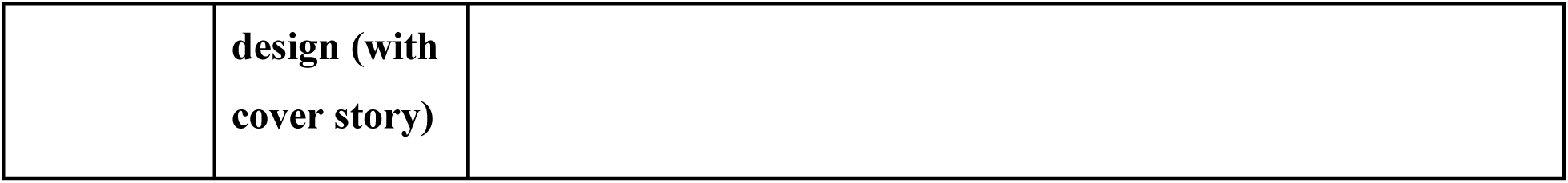
Summary of the experiments

| **Experiment type** |  | **Social detection** | **Social discrimination** |
| --- | --- | --- | --- |
| **Task** |  | Presence vs. absence of social interactions | Discrimination: playful vs. aggressive interactions |
| **Parametrized motion attribute** |  | Chase directness | Charge speed |
| **Sample size after excluding bad participants (see “Exclusion criteria”)** | **Pilot experiments (free-text responses)** | N = 60 (without cover story; 33 females, 27 males; $33.88 \pm 9.18$ years old) | N = 102 (with cover story; 34 females, 59 males; $35.16 \pm 9.59$ years old)<br>N = 103 (without cover story; 47 females, 52 males; $33.71 \pm 10.14$ years old) |
| | **Main experiments (with cover story)** | N = 312, session 1 (149 females, 162 males; $40.70 \pm 12.87$ years old). N = 240 of them returned for session 2 | N = 319, session 1 (138 females, 177 males; $37.43 \pm 12.06$ years old). N = 269 of them returned for session 2. |
| | **Control experiment (invisible “chase” stimuli; with cover story)** | N = 308 (162 females, 144 males; $38.55 \pm 12.75$ years old) | NA |
| | **Within-subject mixed-task** | N = 279 (134 females, 140 males; $36.55 \pm 12.44$ years old) | |

|  |  |
| --- | --- |
|  | **design (with**<br>**cover story)** |

Participants might have been using visual features other than *chase directness* to form their judgments of socialness. For example, past work has shown that agents that are closer together are more likely to be perceived as interacting (Rasmussen & Jiang, 2019). That more-direct chases also resulted in a narrowing of the distance between the two agents over time was an unavoidable consequence of our animation-generation algorithm (where the predator was programmed to chase after the prey); indeed, in our stimulus set, chase directness and mean distance between agents over the course of the animation were correlated across animations (Pearson *r* = –0.69, *p* < .001). Hence, to account for other visual features beyond *chase directness* that participants may have been using, we ran additional models including mean distance between agents as a covariate. While this term was also a significant predictor of socialness ratings (*b* = –0.432, *p* < .001) and model including this term fit the data better (AIC – 12325 compared to AIC –12293 for the model without mean distance; lower AIC indicates a better fit), *chase directness* had a significant effect on the socialness ratings even in the model including mean distance (*b* = 0.603, *p* < .001). Further, mean distance was not a predictor of the predator identification accuracy (*b* = –0.716, *p* = .179) and also did not meaningfully improve the model fit for accuracy (AIC without and with mean distance = –7986 and –7984, respectively).

Lastly, participants may have been relying on other heuristics to form socialness judgments. Since a chase typically results in correlated movement patterns between two agents (when the prey changes direction, the predator is likely to do so a moment later), this correlation will be stronger for more direct than less direct chases. However, agents can also show correlated motion without one necessarily chasing the other—i.e., one agent could change direction every time the other one does but be equally likely to turn away from (or orthogonal to) the path of the other. To test whether participants are simply relying on nonspecific correlated motion as a heuristic, also inspired by Gao et al. (Gao et al., 2009), we ran a separate control experiment in an independent set of participants where we replaced half of the directness 0.167 to 1 chases (6 levels) with a non-social “invisible chase” control. In these trials, the predator was chasing a true prey agent that was made invisible to observers, while a visible “fake” prey mimicked the true prey’s trajectory reflected over a 180° rotation. In this way, correlated motion between the two agents was preserved—when the true prey changed direction, so did both visible agents (the predator and the mimicking agent)—but not necessarily in a manner consistent with chasing. In line with our prediction, we saw that while in the true chase condition, socialness ratings increased with *chase directness* (*b* = 0.471, *p* < .001), in the invisible chase condition, if anything, there was a slight trend in the opposite direction (socialness ratings decreased as directness increased; *b* = –0.146, *p* < .001; supplementary Fig S3b). Both models also included mean distance between agents as a predictor term, and in both conditions, as mean distance increased, the animations were seen as less social (chase: *b* = –0.684, *p* < .001; invisible chase [control]: *b* = –0.493, *p* < .001). This suggests that the increase in socialness with *chase directness* is not merely because of correlated motion in general, but rather correlated motion that is specifically consistent with pursuit behavior. In sum, in line with Gao et al., these experiments show that the motion attribute *chase directness* influences how social stimuli are perceived to be at the group level.

### Discrimination task

Once a social interaction is detected, the next step is to discriminate the *nature* of that interaction; past work (Castelli et al., 2000; Heider & Simmel, 1944) and real-life experience suggest that given the same sensory input, there may be even more variability across people in their percepts of *how* agents are interacting than simply *if* they are interacting. Interactions can be characterized along several dimensions, but a fundamental one is valence—i.e., how positive or negative is the interaction? (Santavirta et al., 2024) Using our parametric approach, we explored changes in the valence of social percepts between "playing" and “fighting”. Playing and fighting, which are preserved throughout much of the animal kingdom, involve two agents moving apart and coming back together in quick succession. We manipulated the speed with which our agents approached each other (*charge speed*) to determine whether this simple motion cue could reliably affect percepts of an interaction’s valence, with slower speeds looking more friendly (like “playing”) and higher speeds looking more aggressive (like “fighting”).

In these experiments (see Table 1), participants watched each animation and rated it on a continuous scale from “playing” to “fighting”. We found that as the *charge speed* increased, interactions were perceived as more aggressive (*b* = 0.622, *p* < .001; Fig 3b). Unlike in the detection task, this relationship, at least within the tested range, was linear, as verified by the comparison between linear and sigmoid curve fits (supplementary Fig S2b; mean difference in model fits, AIC_sigmoid_ – AIC_linear_ ≥ 3.47, *p* < .001 in all the discrimination experiments). The effect of *charge speed* persisted when controlling for the mean distance between the two agents (which also affected ratings such that higher mean distances predicted slightly less aggressive ratings; *b* = –0.057, *p* = .008)) and trial number as an index of time (for which we found that percepts of aggressiveness also weakly increased over the course of the experiment; *b* = 0.024, *p* < .001).

Together, results indicate that social percepts ⎯ both the presence and nature of an interaction ⎯ can be manipulated by simple motion attributes in ways that are generally shared across people. The differences in the shapes of the tuning curves between experiments (sigmoidal for detection, linear for discrimination) may be because presence is more of a categorical variable while valence is a more continuous property, though it is also possible that the discrimination curve would become sigmoidal at more extreme *charge speeds*. To what extent are there stable and meaningful individual differences in these socio-perceptual tendencies? We probe this question next.

### Robust individual differences in social perception exist atop group-level tendencies

Even given these shared general tendencies, individuals often vary in their percepts of social interactions, especially when faced with ambiguous scenarios. To quantify this across-subject variability, inspired by psychophysics approaches, we fit individual participants’ rating data with a sigmoid (detection experiments) or linear function (discrimination experiments; see **Methods**) to derive individual “social tuning curves”. For detection curves, we focused on three main parameters: (i) point of subjective equality (*PSE*), the value of *chase directness* at which the participant’s percept crosses the midpoint of the rating scale (0.5); higher values indicate that more evidence is needed to declare something “social”; (ii) *range*, the difference between ratings at the lowest (0) and highest (1) levels of *chase directness*; higher values may reflect higher perceptual vividness, certainty or diversity; and (iii) *bias*, the extent to which ratings are skewed toward one end of the scale; higher values (> 0.5) indicate a bias toward social (“chasing”) while lower values (< 0.5) indicate a bias toward nonsocial (“moving independently”). For discrimination curves, we focused on two main parameters: (i) *PSE*, the *charge speed* at which participants’ percept crosses the midpoint (0.5; i.e., percept switches from play to fight), lower values reflect a lower threshold to detect aggressive interactions ; and (ii) *bias*, the extent to which ratings are skewed toward one end of the scale; higher values (> 0.5) indicate a bias toward “fighting” while lower values (< 0.5) indicate a bias toward “playing”. See Methods Fig 2 for a schematic of these parameters.

While the vast majority of participants showed the same general directionality as the group-level trends, fine-grained properties of tuning curves differed between participants (see Fig 4a for sample participants [all data in supplementary Fig S4 and S5] and Fig 4b for full distributions of parameters of interest). To test the stability of these tuning curves within participants, we had participants return for a second session 1–2 months later, in which the task design was identical to the first session except that we used previously unseen animations (generated using the same algorithms). Visual inspection showed that idiosyncrasies in behavior and tuning curves were largely preserved across sessions (e.g., Fig 4a). The main curve parameters did show some systematic session-to-session variation in both tasks, but their magnitudes were very small: namely, in the detection task, people discriminate slightly less between social and non-social interactions (lower *range*) in session 2, and in the discrimination task, people are slightly less biased towards aggressive ratings in session 2 (see supplementary Fig S6a for details).

**Fig 4:**
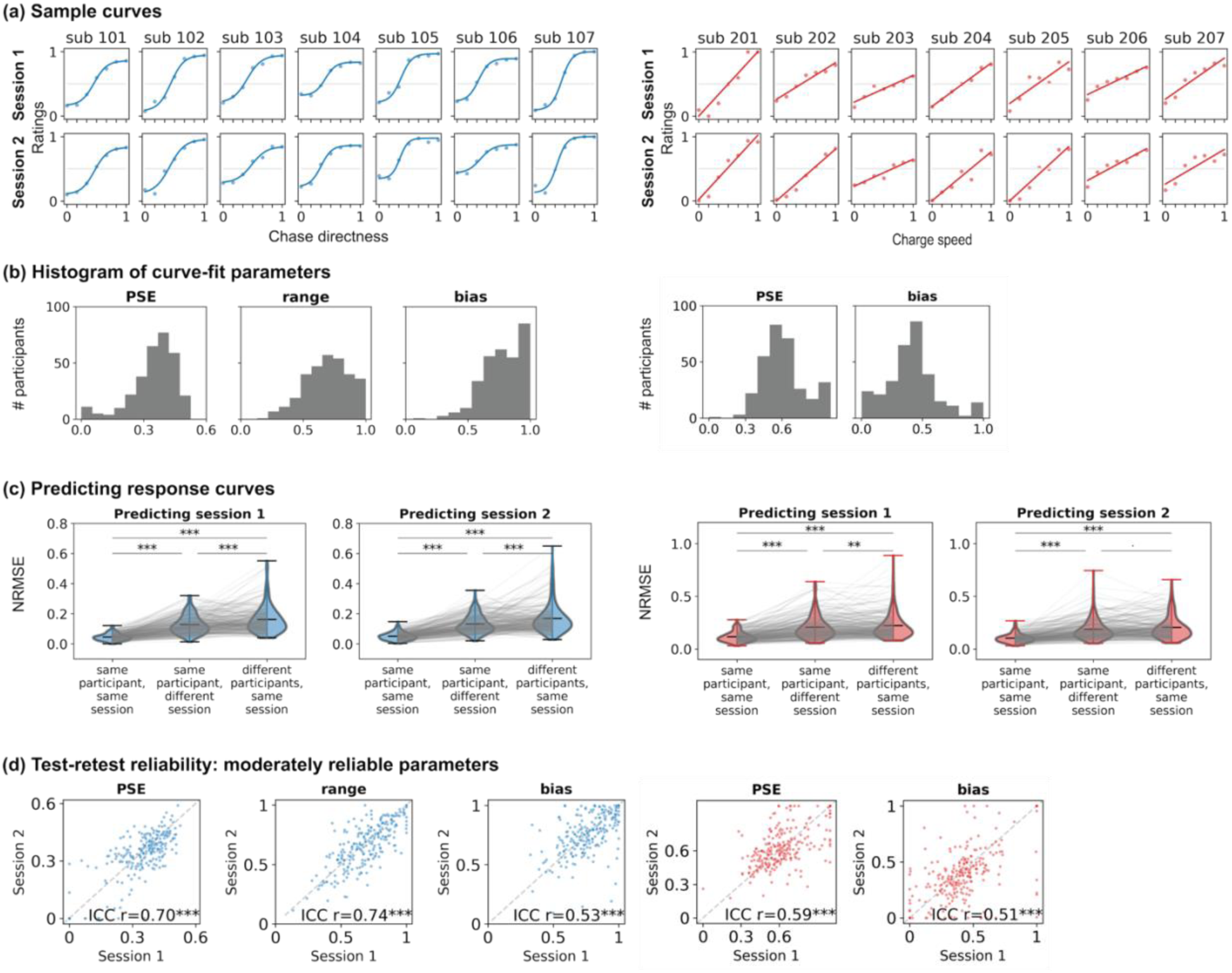
Individual differences in social tuning curves. (a) Tuning curves for sample participants in two distinct sessions. Dots represent mean ratings at each motion attribute level, and the line shows the best fitting sigmoid (detection) or linear curve (discrimination). For the full versions, see supplementary Fig S4 and S5. (b) Histograms showing the full distributions of the main curve parameters of interest from all participants in session 1 (left, detection experiment; right, discrimination experiment). PSE, point of subjective equality. (c) Predicting individuals’ single-session ratings using curve parameters fit to their own data from the same session (left), their own data from a different session (middle), or the average parameters from all other participants in the same session (right). NRMSE=normalized root mean square error; lower values indicate better fits. (d) Test-retest reliability between sessions 1 and 2 of the main curve parameters of interest. Each dot represents a participant. Test-retest reliability for other parameters is shown in supplementary Fig S6. *** = p < .001, ** = p < .01, ^.^ = p < .1.

**Fig 5:**
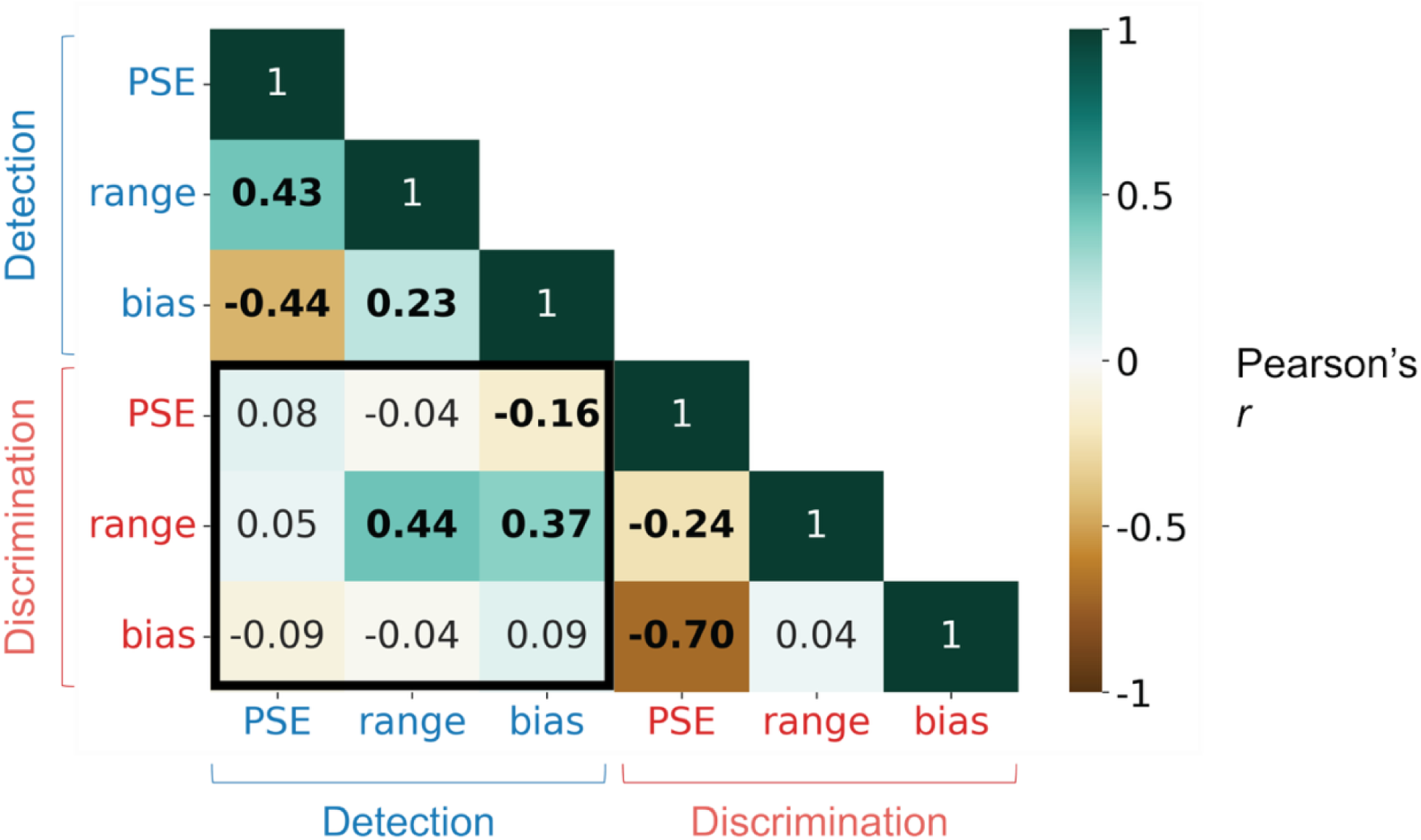
Correlation between the main curve parameters of interest from the social detection and discrimination tasks in the mixed-task experiment (within-subject design). Across tasks (black box), two moderate and one weak pairwise correlation emerged, suggesting that each task provides non-redundant information about individuals’ socio-perceptual tendencies. Significant correlations (FDR q < .05) are displayed in bold text.

We quantified the stability and uniqueness of tuning curves in two ways. First, we used parameters calculated on data from a participant’s first session to fit the same participant’s data in their second session (or vice versa). To test the extent to which curve parameters were both stable within people and distinct across people, we compared this within-participant fit to the mean across-participant fit calculated by using each participant’s curve to fit data from all other participants in the same session. Participants’ ratings were generally better predicted by their own curves from a different session than the average of everyone else’s curves in the same sessions (Fig 4c, middle and right violin plots within each subplot; outliers omitted for clarity; detection task: mean difference (*MD*) = 0.04 and 0.05 when fitting session 2 data to session 1 and vice versa, both *p* < .001; discrimination task: *MD* = 0.07 (*p* = .006) and 0.08 (*p* = .06) when fitting session 2 data to session 1 and vice versa). Second, we calculated the intra-class correlation coefficient between parameters fit to data within each session for each parameter of interest. In the detection task (sigmoid fit), *PSE*, *range* and *bias* showed moderate to good reliability; in the discrimination task (linear fit), *PSE* and *bias* showed generally moderate reliability (Fig 4d; see supplementary Fig S6b-c for data on other parameters that were less reliable and/or redundant with the main curve parameters of interest and supplementary Fig S6d for the covariance between all curve parameters). Overall, then, individual tuning curves for both social detection and discrimination were both stable within participants (reliable across sessions) and unique between participants, suggesting a trait-like component.

### Individual social detection and discrimination tendencies are moderately related

Social detection and discrimination both exhibit shared tendencies as well as robust individual differences, but how do behaviors relate across the two tasks? In other words, are individuals’ discrimination tendencies predictable from their detection tendencies (and vice versa)? For example, people who are more prone to seeing social interactions in the first place might also be more prone to seeing them in a more positive (or negative) light. To study this, we conducted a third experiment in which a new set of participants performed both the detection and discrimination tasks in interleaved blocks in a single session. We successfully replicated the group-level behavioral trends (Fig 3) in this new group of people (see supplementary Fig S8).

We then fit sigmoid and linear curves to each individual’s detection and discrimination data, respectively.

Next, we correlated curve parameters across participants both within and across tasks.

The within-task correlations were overall similar to the correlations from the individual detection and discrimination tasks (supplementary Fig S6d). Focusing on between-task correlations (highlighted part of the matrix in Fig 5), the majority of pairwise correlations were weak, suggesting that detection and discrimination behavior are overall relatively independent. We did, however, find three significant relationships that survived multiple comparison correction (false discovery rate or FDR *q* < .05). First, the *range* parameter correlated moderately across tasks (r = 0.44). This indicates that people who distinguished social (“chasing”) from non-social (“moving independently”) more strongly also distinguished negative interactions (“fighting”) more from positive interactions (“playing”) more strongly. This could reflect people’s general confidence/willingness to use extremes (people who are more confident may have used a wider range of the rating scale in both tasks) and/or the extent to which their percepts are sensitive to sensory evidence (people whose percepts vary more strongly with sensory evidence would have a higher range in both tasks). Next, people who showed a higher bias toward socialness (“chasing”) in the detection task also showed a higher range (more distance between extremes) in the discrimination task (*r* = .37). This suggests that people who are more predisposed to detecting social interactions may also be more sensitive to motion cues and/or more certain when discriminating between different types of interactions. Lastly, people who showed a higher bias towards socialness had a lower PSE, i.e., they characterized interactions as aggressive at lower *charge speeds* (*r* = −0.16). Still, on the whole, social discrimination tendencies were largely not directly predictable from detection tendencies and vice versa. This suggests that behavior on the two tasks may be complementary in revealing individual differences in social perception, and combining information about detection and discrimination tuning curves likely better characterizes individuals’ socio-perceptual tendencies than one task alone.

### Traits explain some of the individual differences in socio-perceptual tendencies

How do socio-perceptual tendencies as measured by behavior on our detection and discrimination tasks relate to real-world variability in social function? In past work, behavior on related tasks has been found to differ in various clinical and subclinical conditions including autism (Klin & Jones, 2006; Lisøy et al., 2022; Rasmussen & Jiang, 2019; Vandewouw et al., 2021; Zwickel et al., 2011), internalizing symptoms (Varrier & Finn, 2022), depression (Kaletsch et al., 2014; Kohler et al., 2011; Krause et al., 2021; Liu et al., 2012) and loneliness (Gardner et al., 2005; Powers et al., 2014). In our final set of analyses, we studied if and how properties of individuals’ social tuning curves covaried with social, affective, and personality traits as measured by established self-report scales.

We performed a series of multiple linear regressions using the trait scores (14 total) to predict two curve parameters each from the detection and discrimination tasks. The first parameter was a combined index, created by subtracting *PSE* from *bias* (*bias – PSE*; see **Methods** for details); this term captures sensitivity to see social and aggressive interactions in the detection and discrimination tasks, respectively. The second was the *range* parameter. We performed separate regressions for the detection and discrimination task (4 regressions in total).

For the detection task, the multiple regression to predict *bias – PSE* showed that individuals with lower communication deficits (*comm*) and agreeableness scores (*agree*) but higher extraversion scores (*extr*) were more likely to perceive social interactions (*95%CI*: *comm* = [–0.89, –0.19], *agree* = [–0.64, –0.12], *extr* = [0.002, 0.78]; Fig 6a, left). In addition to these specific relationships, the full detection model performed well in a permutation test to test its overall exploratory power (where *bias* – *PSE* was permuted 10k times and their *R^2^* values compared against the true model *R^2^*; *p* = .03; Fig 6a, left, inset). For the discrimination task, the multiple regression showed that individuals who were less lonely were more likely to perceive aggressive interactions in the (*95%CI*: *lone* = [–0.83, –0.09], Fig 6a, right); however, the overall discrimination model performed poorly in a permutation test (*p* = .5; Fig 6a, right, inset), suggesting that this result should be interpreted with caution.

**Fig 6:**
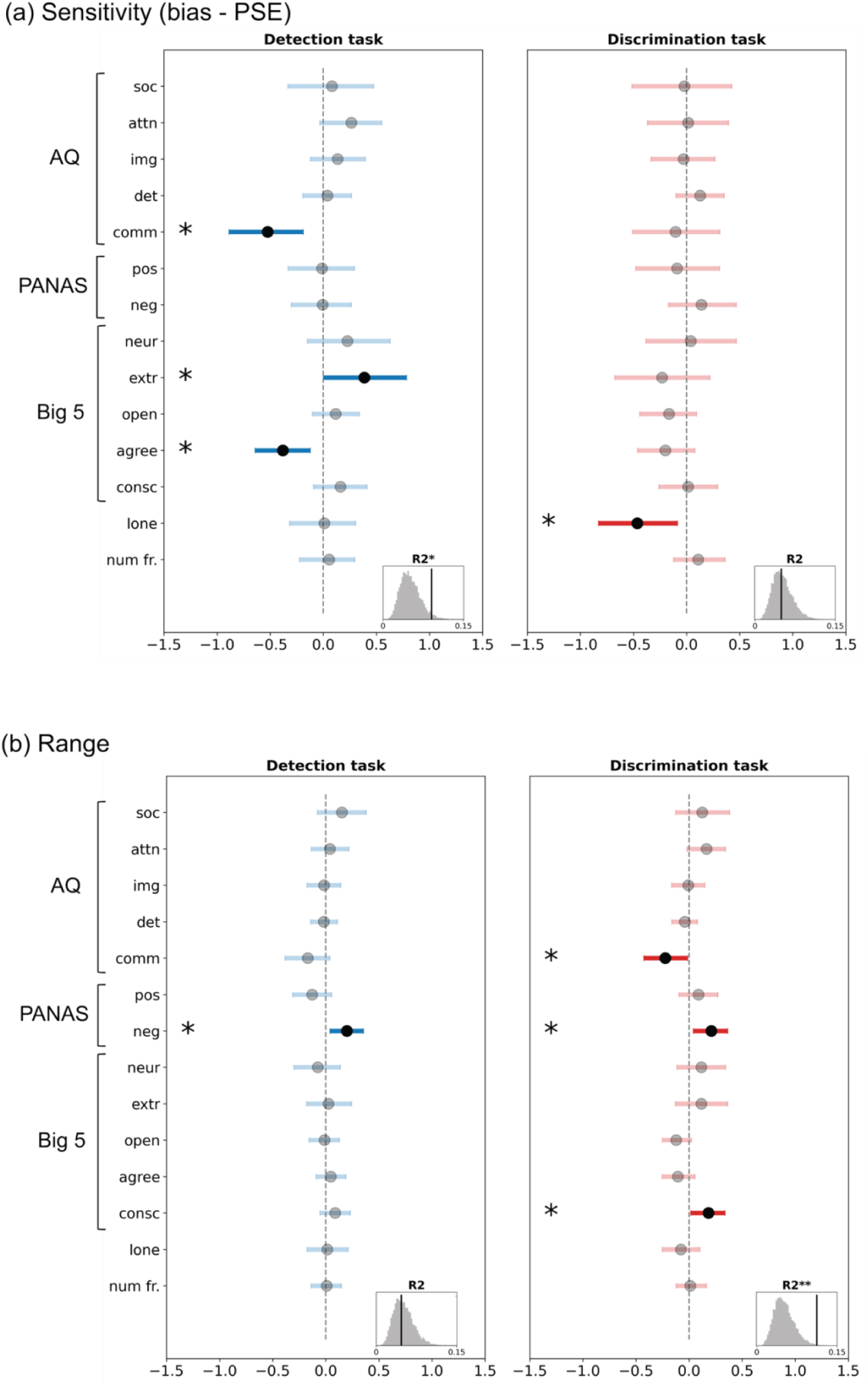
Results from the multiple linear regression between task behavior (curve parameter = f(traits)) and the 14 trait dimensions. Regression estimates for (a) the combined sensitivity index bias – PSE, which reflects sensitivity towards social (detection) or aggressive (discrimination) interactions and (b) range. The statistical significance of the regression coefficients was obtained by bootstrapping (sampling with replacement) participants 1000 times and estimating the 95% CI of each regression coefficient. The black markers show the mean, and the blue/red lines show the 95% CI. The histograms at the lower right section within each subplot show how the true model fit (vertical black line) compares to 10,000 permutations (shuffles of the curve parameter; grey histogram). *= p < .05, ** = p < .01, ***= p< .001. Abbreviations: Autism Quotient (AQ) questionnaire subscales – ’soc’: social skill deficits, ’attn’: attention-switching deficits, ’img’: imagination deficits, ’det’: heightened attention-to-detail and ’comm’: communication deficits; Positive and Negative Affect Schedule (PANAS) subscales – ‘pos’: positive affect and ’neg’: negative affect; Big 5 (NEO-FFI) subscales – ’neur’: neuroticism, ’extr’: extraversion, ’open’: openness, ’agree’: agreeableness and ’consc’: conscientiousness; ‘lone’: loneliness score; ‘# fr.’: number of friends.

The multiple regression to predict *range* showed that individuals with higher negative affect (*neg*) had a higher *range* (used more of the non-social to social scale) in the detection task (*95%CI*: *neg* = [0.04, 0.35]; Fig 6b, left) and that people with lower communication deficits, higher negative affect and conscientiousness had a higher *range* in the discrimination task (used more of the play-fight scale; *95%CI*: *comm* = [–0.43, –0.01], *neg* = [0.04, 0.36], *C* = [0.02, 0.34]; Fig 6b, right ). Here the overall goodness of model fit was the opposite of what we saw with *bias – PSE*; while the discrimination model fit was better than the permutations (*p* = .004; Fig 6b, right, inset), the detection model performed poorly (*p* = .55; Fig 6b, left, inset).

To conclude, we found some evidence that individual differences in social sensitivity (*bias – PSE* from the detection task) and the discriminability between playful and aggressive interactions (*range* from the discrimination task) showed some relationships with socio-affective traits. In particular, the relationships with social sensitivity in the detection task are in line with past work and our *a priori* hypotheses that individuals who are more skilled at social interactions (lower communication deficits) and more motivated to engage in social interactions (higher extraversion) are also more likely to perceive social information in their environment. The relationships with *range* in the discrimination task should be interpreted in light of our cover story, which induced a general expectation of playfulness; the significant terms suggest that higher general negative affect, more carefulness (higher conscientiousness) and stronger social skills (lower communication deficits) may underlie better discriminability between positive and negative social interactions, or, in other words, less of a bias to see only playful interactions.

## Discussion

Here, we used a psychophysics-inspired approach to characterize both group-level tendencies and individual differences in social perception. We found strong commonalities in how people use relatively low-level motion attributes to arrive at percepts of the presence (detection experiments) and nature (discrimination experiments) of a social interaction, and also robust individual differences that were replicable over a period of months and showed some relationships to trait phenotypes.

Our approach lends a level of rigor and precision to the study of social perception. While some notable past work has used parametric stimulus manipulations (Gao et al., 2009; Horovitz et al., 2004; Johansson et al., 2004; Looser & Wheatley, 2010; Schultz & Bülthoff, 2019), here, we extend this approach to individual-subject data to recover single-person social tuning curves that are both reliable and unique. Generating stimuli algorithmically makes our approach simultaneously more objective and more subjective: more objective because we can create parametric manipulations using quantitatively defined features (rather than relying on handcrafted stimuli created and labeled via experimenter intuition (Castelli et al., 2000)) and thereby generalize beyond item-level effects, and more subjective because we are eschewing any notions of a ground truth and classifying behavior according to observers’ own reports (Peters, 2024), which better reflects what happens in the real world—where different people can and do interpret the same social situation differently.

Social cognition is traditionally considered a high-level process, but more recent work has found evidence that recognizing and processing social information begins earlier in the perceptual hierarchy than previously thought (Gandolfo et al., 2024; Malik & Isik, 2023; McMahon & Isik, 2023; Pitcher & Ungerleider, 2021; Thieu et al., 2024), and artificial intelligence can extract basic cues as to the presence and nature of social information using fast, automatic, visually-based processes (Malik & Isik, 2023). The recently proposed “third visual pathway” in the brain, which runs along the lateral surface from early visual regions into the superior temporal sulcus and is specialized for extracting social information from dynamic cues, embodies the theory that our visual system might be especially attuned to social information, given its evolutionary importance (Pitcher & Ungerleider, 2021). Our work adds to this growing body of work by showing that parametrically varying meaningful “mid-level” visual motion attributes (McMahon et al., 2023) even in very stripped-down, simplified stimuli can directly modulate social percepts at both the group and individual level, thus confirming that social perception involves visual evidence accumulation with both an objective and subjective component. Of note, autism is a condition marked by deficits with social cognition, but also altered basic visual processing (Choi et al., 2023; Robertson & Baron-Cohen, 2017); the idea that perception is the fundamental starting point for social cognition might lead us to discover hierarchical links between aberrations in these two domains.

While the motion attributes we used here, *chase directness* and *charge speed*, were sufficient to evoke varying social detection and discrimination percepts respectively, we also acknowledge that social percepts are governed by many more dimensions than these two and that the cover story may have restricted the possible range of interpretations. We propose the idea of individual social perception “landscapes” that can be conceptualized as a multidimensional space spanned by objectively defined axes (e.g., motion attributes such as our *chase directness* and *charge speed*, plus many others) where the dimensions themselves are fixed, but sensitivity to these dimensions varies across people and can be expressed in terms of tuning curve parameters.

In Fig 7, we show a two-dimensional schematic of what these landscapes might look like and how they might vary. In this example, the horizontal axis represents an attribute that influences *if* a social interaction is perceived (detection; e.g., *chase directness*) and the vertical axis represents an attribute that influences *how* a social interaction is perceived (discrimination; e.g., *charge speed*). A healthy neurotypical individual (Fig 7a) might show moderate sensitivity to objective evidence for socialness (indicated by the saturation gradient along the horizontal axis); once information is generally deemed social, percepts of the *nature* of that information are generally balanced between positive and negative interactions (even vertical distribution of pink and blue). Someone with an autism-like phenotype (Fig 7b) might have lower sensitivity to social information—in other words, they might require higher doses of objective evidence to detect a social interaction. For someone with a depression-like phenotype (Fig 7c), detection sensitivity may be largely normal, but discrimination might be skewed toward negative percepts. Lastly, someone with psychosis/paranoia-like traits (Fig 7d) might show both heightened sensitivity to socialness and a bias towards negative percepts—in other words, a proneness to read social intentions, particularly nefarious ones, into scenarios that others might perceive as non-social or, at most, social yet neutral. While we explored only two possible dimensions (in separate experiments and blocks) here, and did not include clinical populations, we see the present set of experiments as a first step toward discovering and characterizing these landscapes—which, because they are based on more implicit behavioral readouts, are possibly less prone to overt bias than self-report measures and therefore a useful complement to existing trait scales. In support of this, our results showed that each axis carries some unique variance in its association with socio-affective traits – while both social sensitivity in the detection task and range in the discrimination task were associated with better communication skills, social sensitivity was also uniquely associated with higher extraversion and lower agreeableness, while discrimination range was associated with higher negative affect and conscientiousness. Adding more dimensions (i.e., using more complex, yet still parameterized stimuli) will likely enhance our ability to characterize real-world social and affective function.

**Fig 7.**
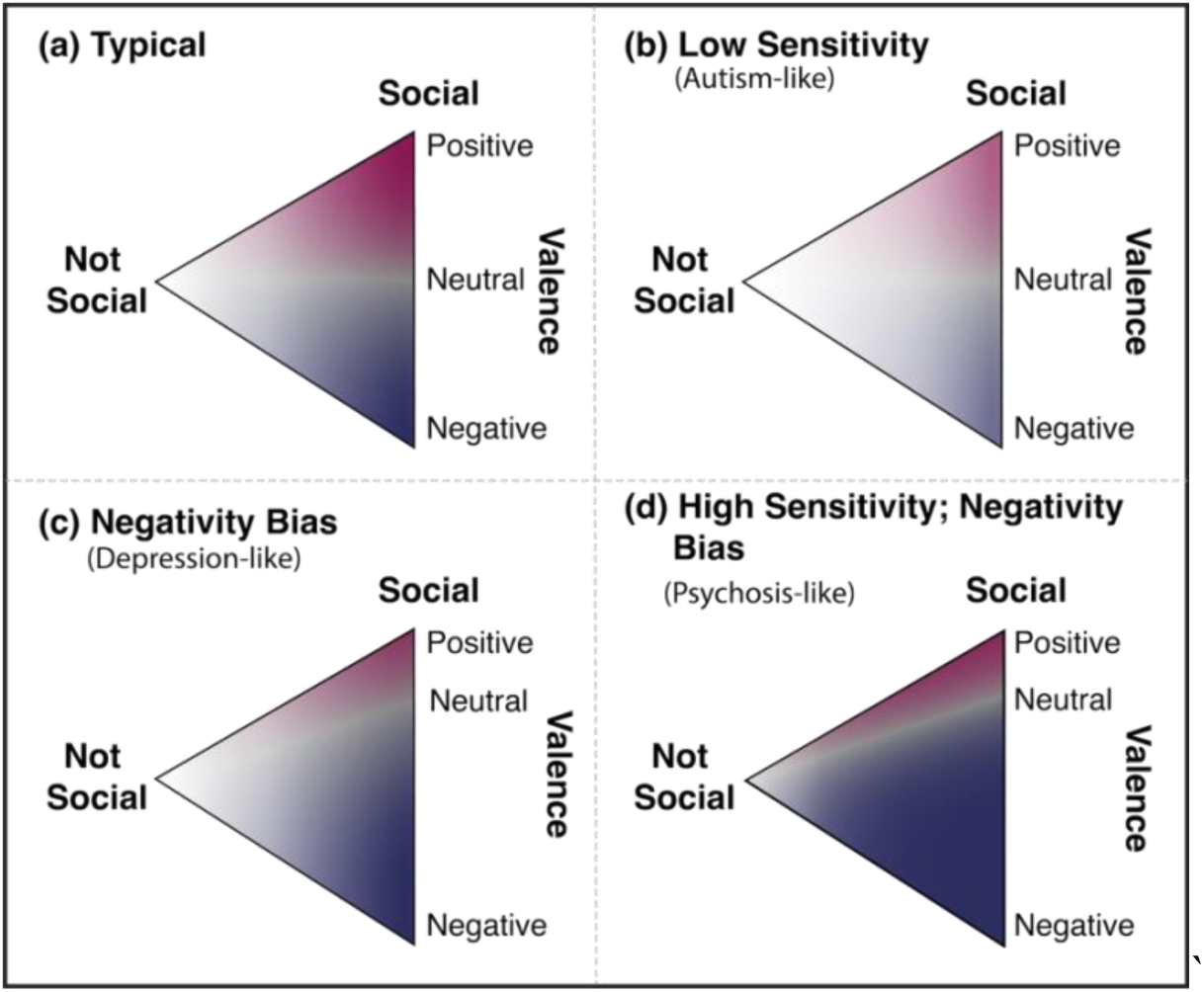
2D projection of social perception landscapes with detection (not social ⟷ social) and discrimination (positive ⟷ negative) along the horizontal and vertical axes, respectively. How subjective percepts of the presence and nature of social interactions may present in (a) a neurotypical individual versus individuals with (b) autistic, (c) depressive or (d) psychotic traits who show, respectively, reduced progression towards social with sensory evidence (large faded area), increased biases towards negative percepts (large blue zone even at typically neutral or positive evidence), and an increased social sensitivity as well as bias towards negative (leftward saturation combined with expanded blue territory), respectively.

Future work can combine our psychophysics-inspired task framework with additional readouts such as reaction times, eye-tracking, physiological measures and/or neuroimaging to yield a more comprehensive picture of the evidence accumulation and decision-making strategies, attentional processes, and other computations underlying individuals’ social-perceptual judgments. There may be a role for generative AI in bridging the gap between the highly impoverished stimuli used here and the full complexity of real-world social information—i.e., we may be able to use generative AI to create more naturalistic-feeling stimuli that are nevertheless still parameterized along known axes. Future work can also tease apart the role of animacy in evoking social percepts by additionally collecting animacy ratings and/or leveraging additional attributes (e.g., facingness, motion jitter) that ensure an even animacy across parametric levels of a motion attribute. In closing, we note that while the vast majority of past work on social perception and cognition has focused on passive (third person) perception of others’ interactions—which is indeed an important part of social cognition—many of our most salient and important social experiences are ones in which we are an active (first-person) participant. One final advantage of our framework is that it can be easily adapted to a first-person context, in which participants themselves are controlling one of the agents and the other agents are programmed to behave in a certain way toward them. This opens the door to generating and comparing social tuning curves between passive and active scenarios, as well as extracting more latent behavioral readouts such as participants’ movement trajectories, which we anticipate will provide an even richer and more useful picture of social perception at both the group and individual levels.

## Acknowledgements

Withheld for anonymity.

