## Supplementary material for "Shared and individual tuning curves for social perception"

For the manuscript:

### Supplementary Methods

#### Stimuli

##### Chase animations

**Chase directness**. Every 350ms (set by an attribute called *subtlety lag* explained in the next paragraph), the predator picks a value from a uniform distribution that ranges from [–*chaseSubtlety*, *chaseSubtlety*], where *chaseSubtlety* ∈ {0°, 30°, 60°, 90°, 120°,150°}. (i.e., *chase directness* ∈ {1, 0.833, 0.667, 0.5, 0.333, 0.167}). These behaviors are illustrated in Fig 1a. To get a feel for what these values mean, we encourage readers to watch some of the animations, available on GitHub. The stimuli used for session 1 chase detection experiments here (https://github.com/thefinnlab/psyanim_behav_paper1/tree/master/stimuli/detection_subtlety/Chase), and those for session 2 experiments are here (https://github.com/thefinnlab/psyanim_behav_paper1/tree/master/stimuli/detection_subtlety/Chase_set2).

**Other attributes**. All attributes besides *chase directness* were set to be constant across animations. Some of the relevant attributes that influenced how the animations were perceived were optimized by manual piloting during stimulus development. These were, for the predator: (i) *maximum chase speed* = 1.5px/frame (frame rate = 60), (ii) *maximum chase acceleration* = 0.1px/frame$^2$ and (iii) *subtlety lag* = 350ms (how often the agent recomputes its chase direction; lower values will make the animation more jittery). For the prey: (i) *flee subtlety* = 30° (the angle which an agent can deviate from the true direction away from the predator — this parameter helps to avoid fleeing looking very obvious), (ii) *safety distance* = 100px (distance below which the agent flees from the predator; above this, it wanders), (iii) *maximum flee speed* =

1.8px/frame, (iv) *maximum flee acceleration*= 0.15px/frame$^2$, (v) *maximum wander speed* = 1.5px/frame, (vi) *maximum wander acceleration* = 0.1px/frame$^2$, (vii) *maximum seek speed* = 3.5px/frame and (viii) *maximum seek acceleration* = 0.05px/frame$^2$ (the seek behavior is included in the prey algorithm to keep it away from the boundaries/walls/edges of the world so that it does not get stuck in a corner). The flee speed and acceleration of the prey were set to be slightly higher than those of the predator to ensure that the predator never actually catches the prey.

#### Wander behavior

We also generated animations where both the agents were wandering independently, meaning that there was no programmed contingency between their motion patterns. The speed and acceleration of the two wandering agents matched that of the predator and prey agents in the main chase animations (pseudo-predator and pseudo-prey agents, respectively) to make these animations as close as possible to the chase animations. These animations serve as an additional check as to whether participants use the speed of an agent as a heuristic to help identify the predator (since, in chase animations, the prey always moved slightly faster than the predator to avoid being caught as described above). Although these animations were generated differently than the chase animations described above (where one of the agents is designed to chase the other, however inefficiently), they are conceptually equivalent to a chase with *chase directness* = 0 (subtlety 180°; i.e., where the predator is equally likely to move in any direction irrespective of the prey's position). Hence, we use these as our experimental stimuli for *chase directness* = 0 in the social detection experiments. These stimuli can be seen here (https://github.com/thefinnlab/psyanim_behav_paper1/tree/master/stimuli/detection_subtlety/Wander/180).

The relevant parameters for these animations in *psyanim* were: (i) *maximum wander speed* (for the pseudo-predator agent = 1.5px/frame,  pseudo-prey agent = 1.8px/frame) , (ii) *maximum wander acceleration* (for the pseudo-predator agent =  0.1px/frame$^2$, for the pseudo-prey agent = 0.15px/frame$^2$), (iii) *maximum angle change per frame* = 35° (how much the movement direction can change from one frame to the next), (iv) *minimum screen boundary distance* = 50px (the distance agents try to maintain from the screen boundary).

#### Invisible "chase" control

There were only 2 relevant parameters for the mimicking agent in *psyanim*: (i) *name or ID of the agent* to mimic (i.e., the invisible true prey), (ii) *angleOffset* = 180° (how much to offset the movement in angles).

#### Discrimination task

As with the detection task, several features were kept at constant values across levels of *charge speed* after extensive in-lab piloting. These were: (i) *minTargetDistanceForCharg*e = 200px, *maxTargetDistanceForCharge* = 500px (the between-agent distance range within which collisions would be initiated), (ii) *mean charge delay* = 200ms, *jitter* = 100ms (how long the second agent waits after the first agent charges at it), (iii) *mean break duration* = 2000ms, *jitter* = 200ms (the duration for which two agents wander between consecutive charges), (iv) *maximum wander speed* = 1.5px/frame, (v) *maximum wander acceleration* = 0.1px/frame$^2$, (vi) *wander panic distance* = 800px (minimum distance at which a wandering agent will charge back at another agent).

#### Quality checks

We quality checked all generated animations for both the detection and the discrimination tasks. Animations were manually checked by at least two lab members and were removed if they contained glitchy/flickery movement patterns, if the agents got stuck in the corners or stuck to each other for extended periods, and/or if one or both agents went offscreen. At all stages, bad animations were replaced with new ones of the same type (e.g., same *chase directness* value, predator color and start position) to obtain the target number of animations. For the detection task, our final stimulus set included 84 chase animations (12 animations at each of 7 *chase directness* levels, including animations generated via the wander algorithm which served as *chase directness* = 0) and 84 invisible "chase" animations (control; 12 animations at each of the *chase directness* levels used in the chase animations plus animations generated via the wander algorithm which served as *chase directness* = 0). Within these sets, the starting position (left versus right of center) and the role of predator versus prey was counterbalanced between the gray and black agent across animations, so that when participants were presented with an animation, there was no expectation of what they would see based on the position or the color of the predator.

For *charge speed* (discrimination) stimuli, we removed bad animations according to similar criteria as described for the detection stimuli. In addition, we ensured that all animations in the final set had the same number of actual collisions (two), since differences in the number of collisions could have influenced percepts independently of *charge speed*, which was the main motion attribute of interest. The final set of discrimination stimuli contained 20 animations at each of 7 *charge speed* levels for a total of 140 animations. Within this set, the initial position of the gray and black agents was counterbalanced across animations.

#### Pilot experiments (open-ended responses)

##### Pilot experiment design

We first conducted a set of small-scale studies where participants could freely describe the animations. Before imposing explicit, constrained rating scales, our goal was to verify that these animations do in fact spontaneously evoke percepts that fall approximately along the intended axes from non-social to social (detection task) or playful to aggressive (discrimination task).

In these experiments, each participant was presented with 7 animations (1 animation for each level of the motion attribute as described under **Stimuli** in the main text). For the detection task, we did not use the cover story. For the discrimination task, we ran two versions – one without and one with the cover story. In what follows, unless otherwise noted, we present data from the version *with* the cover story. After watching each animation, participants responded to the following prompt to indicate what the dots could have represented: "Briefly describe what the dots were doing. Guess if you do not know." (In the discrimination task with the cover story, the text varied slightly: "Describe what the dots were doing using a word or a short phrase"). The task lasted ~5-10 min overall.

##### Pilot experiment data analysis

We analyzed the free-response data using techniques from natural language processing. In each experiment (detection and discrimination tasks), we derived the average "meaning" of descriptions at each stimulus level using semantic embeddings. Specifically, we used Bidirectional Encoder Representations from Transformers (BERT)(Wang et al., 2020) language models as implemented in the Python library *SentenceTransformers* (https://huggingface.co/sentence-transformers/all-MiniLM-L6-v2). For each description (participant response, ranging from a phrase to a short sentence or two), we get a single 384-dimensional vector embedding. We then averaged across all embeddings at each stimulus level (before averaging, detection task: 12 unique stimuli per level of *chase directness* x 5 observers per stimulus = 60 observations per level of *chase directness*; discrimination task: 20 stimuli per level of *charge speed* x 5 observers per stimulus = 100 observations per level of *charge speed*. After averaging, there was 1 embedding per *chase directness* level or *charge speed* level).

In an initial exploratory/data-driven analysis, we compared this mean vector to the embeddings of *all* 8432 English verbs from the natural language toolkit (nltk; https://www.nltk.org/howto/wordnet.html) to identify the 5 verbs whose embeddings it was closest to. To more clearly isolate the *differences* in percepts across levels, we then removed words that appeared in at least 6 of the 7 motion attribute levels within each experiment.

In a follow-up, more hypothesis-driven analysis, we quantified the change in percepts across stimulus levels by computing the similarity of mean embeddings at each level to our expected percepts at either ends of the response scale (detection task: “chasing”, “moving independently”; discrimination task: “playing”, “fighting”). We expected that, as the motion attribute value increased, descriptions’ similarity to one extreme (“chasing” or “fighting”) would increase and similarity to the other extreme (“moving independently” or “playing”) would decrease. To quantify this, we took the difference between embeddings’ similarity scores to both extrema ($difference\_score = score\_chasing - score\_moving\_independently$ for the detection task and $difference\_score = score\_fighting - score\_playing$ for the discrimination task), giving us one $difference\_score$ per trial. Later, scores were compared using a linear mixed effects model (LME; *pymer4* package (Jolly, 2018)): $difference\_score \sim motion\_attribute + (1|sub\_id)$ , where for the detection and discrimination tasks, $motion\_attribute$ referred to *chase directness* and *charge speed*, respectively.

Finally, we performed two additional analyses specific to for the discrimination task. In the first analysis, we quantified how the valence, or "sentiment", of the descriptions varied across attribute (*charge speed*) levels. We used a RoBERTa-base model (Loureiro et al., 2022) (https://huggingface.co/cardiffnlp/twitter-roberta-base-sentiment-latest/tree/main) to automatically quantify sentiment, which yields a positive, negative, and neutral score for each description. Our dependent variable $sentiment\_score$ was the difference between the negative and positive sentiment scores for each description. Similar to the approach to semantic similarity described in the previous analysis, the effect of the motion attribute (namely, *charge speed*) on these values was computed using the LME: $sentiment\_score \sim charge\ speed\ + (1|sub\_id)\ +\ (1|stim\_id)$. In the second analysis, we quantified how arousal varied across attribute levels. For this, we used the National Research Council Canada (NRC) Valence, Arousal and Dominance (NRC-VAD) Lexicon (Mohammad, 2025) which contains arousal scores for individual words. We split each response into words, computed their arousal scores and averaged these scores for each text response to get a single arousal_score. Next, we used the LME $arousal\_score \sim charge\ speed\ +\ (1|sub\_id)\ +\ (1|stim\_id)$ to assess how arousal changes with *charge speed*, and lastly, we ran the model $sentiment\_score \sim charge\ speed\ + arousal\ score\ + (1|sub\_id)\ +\ (stim\_id)$ to assess if changes in arousal explained away effects of *charge speed* on valence.

We noted that free-text descriptions were overall biased toward positive sentiment (Fig S1c, left). This is likely because of our cover story about the dots representing children in a park, which induces a strong prior toward playful interactions. To check this, we ran an additional small pilot batch without a cover story (all else remained the same).

Results from the analyses of free text responses in these pilot experiments showed evidence favoring our hypotheses that (1) in the detection study, as *chase directness* increases, animations are seen as more social, and (2) in the discrimination study, as *charge speed* increases, interactions are seen as more aggressive, with a higher arousal and negatively valenced. Together, the pilot experiments confirmed that the stimuli we generated algorithmically could spontaneously—i.e., without prompting with explicit choices of possible interactions—evoke percepts along the intended axes, and this gave us the confidence to move forward with our main experiments using these axes to structure responses, described in the next section.

#### Main experiments

##### Quality checks during data acquisition

We performed a few data quality checks during data acquisition to exclude poor participants within the first few minutes of the study. First, after the instructions, including the cover story about the dots representing children in a park, but before the start of the main experiment, we presented participants with a multiple-choice question as to what the dots represented. The options were “animals”, “balls”, “adults”, “children”, “magnets”. The correct answer was “children” (as mentioned clearly in the cover story). If participants responded incorrectly, they were given one chance to correct their answer, and if their second response was also incorrect, they were immediately excluded from the study. (Note that this same question was asked again *after* the main task and used for a second quality-check analysis, see below.) Participants were also warned that they may not be compensated if they missed (i.e., timed out on) more than 10%

of trials. We also excluded participants who opened other tabs or had bad internet connections by including a demo animation on the very first page and checking playback duration in real time. If the duration of this page was much higher than 6 or 8s (actual duration of the animations), this meant that the animation did not play as normal, and either paused (because of other open tabs and the participant not paying attention) or played very slowly (possibly because of a slow internet connection). Participants who stayed on the demo animation page for longer than a liberal threshold of 20s were immediately excluded from the study. Besides these online quality checks *during* the experiments, we conducted further quality checks at the data analysis stage to exclude participants with poor data quality.

#### Data exclusion criteria

First, we excluded participants with bad or unreliable data, as defined by meeting one or more of the following criteria: (i) missing responses (i.e., timing out) on more than 5% of all trials; (ii) incorrect responses in the post-main-experiment debrief question asking them to identify what the dots represented (note that this question is identical to the question asked at the beginning of the main experiment, but the rationale here is that if by the end of the main experiment participants had forgotten what the dots represented, their perception and ratings could have been affected by whatever they assumed the dots to represent by the end); (iii) (for the detection task alone) incorrectly identifying the predator in more than one third of animations with *chase directness* = 1 (rationale: the chase/predator identity is very obvious in these animations, so incorrect answers here are most likely failures of attention); (iv) lingering on the animation page for more than 20s in at least 5% of trials (each animation was only programmed to last 6 or 8s, and so the page should have lasted for a similar duration; any longer indicates that they may have clicked away from the experiment tab and/or had a slow internet connection); (v) failing to respond to ≥ 10% of items on one or more trait questionnaires; or (vi) missing at least one (out of five) attention-check items in the trait questionnaires. Trait questionnaires are described in detail in the **Trait measures** section.

#### Trait measures

The AQ consists of 50 total items measuring five subdomains: social skill deficits, communication deficits, attention-switching deficits, heightened attention to details and imagination deficits. For each item, participants had four response choices ("Definitely disagree", "Slightly disagree", "Slightly agree", "Definitely agree"). We reverse-scored the items that were intended to be as per the instructions from the creators; however, when assigning a score on each item, we assigned responses scores between 0 and 3 (where 3 → less neurotypical and more autistic) in place of binarizing responses (assigning 0 to the first two levels and 1 to the last two levels). Higher scores on each subdomain indicate more autism-like traits (e.g., greater social skill deficits).

PANAS consists of 20 total items, 10 measuring positive affect and 10 measuring negative affect. Participants responded on a five-point scale ("Very slightly or not at all", "A little", "Moderately", "Quite a bit", "Extremely"). Each response was coded between 0 and 4, and there were no reverse-scored items. This scale results in separate scores for positive and negative affect.

NEO-FFI consists of 60 total items measuring five dimensions: openness, extraversion, neuroticism, conscientiousness, agreeableness. Participants responded on a five-point scale ("Strongly disagree", "Disagree", "Neutral", "Agree", "Strongly agree") coded between 1 and 5. This scale yields a summary score for each of the five dimensions.

The UCLA loneliness scale consists of 20 total items that measure a single dimension. Participants responded on a 4-point scale ("Never", "Rarely", "Sometimes", "Always") scored between 1 and 4. Higher scores indicate higher loneliness. Lastly, participants also responded to the following question with an integer value: "*Please estimate the number of* ***close friends*** *that you have, where "close friends" are people that you feel at ease with and can talk to about private matters.*"

Within each questionnaire we also added one attention check question (e.g., for AQ, the question was this: "If you are doing your best to complete this survey honestly, choose 'Definitely agree'.") to confirm that participants were paying attention to the questions. The accuracy on these questions was used as a quality check criterion during data pre-processing (described under *Main experiment analysis above*).

### Supplementary Results

#### Animations that vary parametrically along simple motion attributes spontaneously evoke varying social percepts

In the pilot experiments, participants (N = 60 and 102 in the detection and discrimination tasks, respectively) watched 7 animations (one per *chase directness* [6s each] and *charge speed* level [8s each]) and generated free-response text descriptions of each animation.

We quantified responses using tools from natural language processing. First, we computed the average semantic embedding across all responses at each stimulus level for the detection and discrimination studies separately and determined the English verbs that were closest to the average description in semantic space. In the detection task, the average description was closest to mechanical verbs (e.g. "move", "reposition", "lateralize") for stimuli at lower *chase directness* (intended less-social conditions), while the average description was closer to verbs indicating a pursuit (e.g., “follow”, “pursue”) for stimuli at higher *chase directness* (intended more-social conditions). In the discrimination task, the average description was closest to gentler, friendlier words (e.g., “play”, “game”, “dance”) at lower *charge speeds* (intended play-like conditions) and to more aggressive words (e.g., “fight”, “battle”, “combat”) at higher *charge speeds* (intended fight-like condition). Details are in Fig S1a.

To quantify these trends, we directly evaluated semantic similarity of average descriptions at each motion attribute level to the intended percepts at the extrema of each continuum (i.e., similarity to “chasing” relative to “moving independently” for the detection task; similarity to “playing” relative to “fighting” for the discrimination task). (Fig S1b). We see that as *chase directness* (intended socialness) increases, similarity to “chasing” relative to “moving independently” increases ($b = 0.176$, $p < .001$). Similarly, as *charge speed* (intended aggressiveness) increases, similarity to “fighting” relative to “playing” increases ($b = 0.12$, $p < .001$). In both cases, we see that the similarity is not perfectly centered – in the detection task, the similarity to chasing is overall higher than the similarity to moving independently, and in the discrimination task, the similarity to playing is overall higher than the similarity to fighting.

Lastly, for the discrimination task only, we quantified how the valence and arousal of descriptions changed with *charge speed* using automated sentiment analysis. The results from the valence analyses confirmed that the descriptions became more negative as *charge speed* increased ($b = 0.067$, $p = .004$, Fig S1c left) but were still overall positive even at the highest speed. This positive bias was mitigated in the second batch without the cover story albeit with a slightly steeper change in the negative sentiment scores with *charge speed* ($b = 0.128$, $p < .001$, Fig S1c right). The results from the arousal analyses revealed that arousal increased with *charge* *speed* with or without the cover story ($b = 0.188$, $p < .001$ and $b = 0.161$, $p < .001$, respectively). Although both arousal and valence (positive – negative score) showed linear effects with *charge speed*, these were only weakly correlated (Pearson $r = .27$, $p < .001$ and $r = –.08$, $p = .05$). Further, even on adding arousal scores to the linear mixed effects model, the effect of *charge speed* on valence persisted ($b = –0.097$ and –0.129 for the batches with and without the cover story respectively, both $p < .001$). In sum, the relative effect of *charge speed* on response sentiment remained largely the same between the versions with and without the cover story and with or without factoring in the effects of arousal.

Taken together, these results confirmed that percepts of our animations did vary along the expected continuum for each experiment type (chasing versus moving independently for the detection task, fighting versus playing for the discrimination task), which gave us confidence to move forward to our main experiments, in which we replaced free responses with these continua as predetermined rating scales.

### Supplementary figures

**(a) Unique verbs closest to responses at each stimulus level**

**Detection task**

| Chase directness | Most representative verbs |
|---|---|
| 0.00 | move, reposition, circularise, lateralize |
| 0.17 | move, wiggle, lateralize |
| 0.33 | move, reposition, lateralize |
| 0.50 | move, lateralize, reposition |
| 0.67 | follow, forward, pursue |
| 0.83 | follow, move, forward |
| 1.00 | follow, pursue, forward |

**Discrimination task**

| Charge speed | Most representative verbs |
|---|---|
| 1.50 | play, dance, game, sport, mingle |
| 2.75 | play, game, outplay, sport, misplay |
| 4.00 | play, game, sport, outplay, maneuver |
| 5.25 | fight, combat, battle, play, game |
| 6.50 | fight, sport, combat, battle, game |
| 7.75 | play, fight, game, battle, combat |
| 9.00 | fight, battle, fistfight, combat, play |

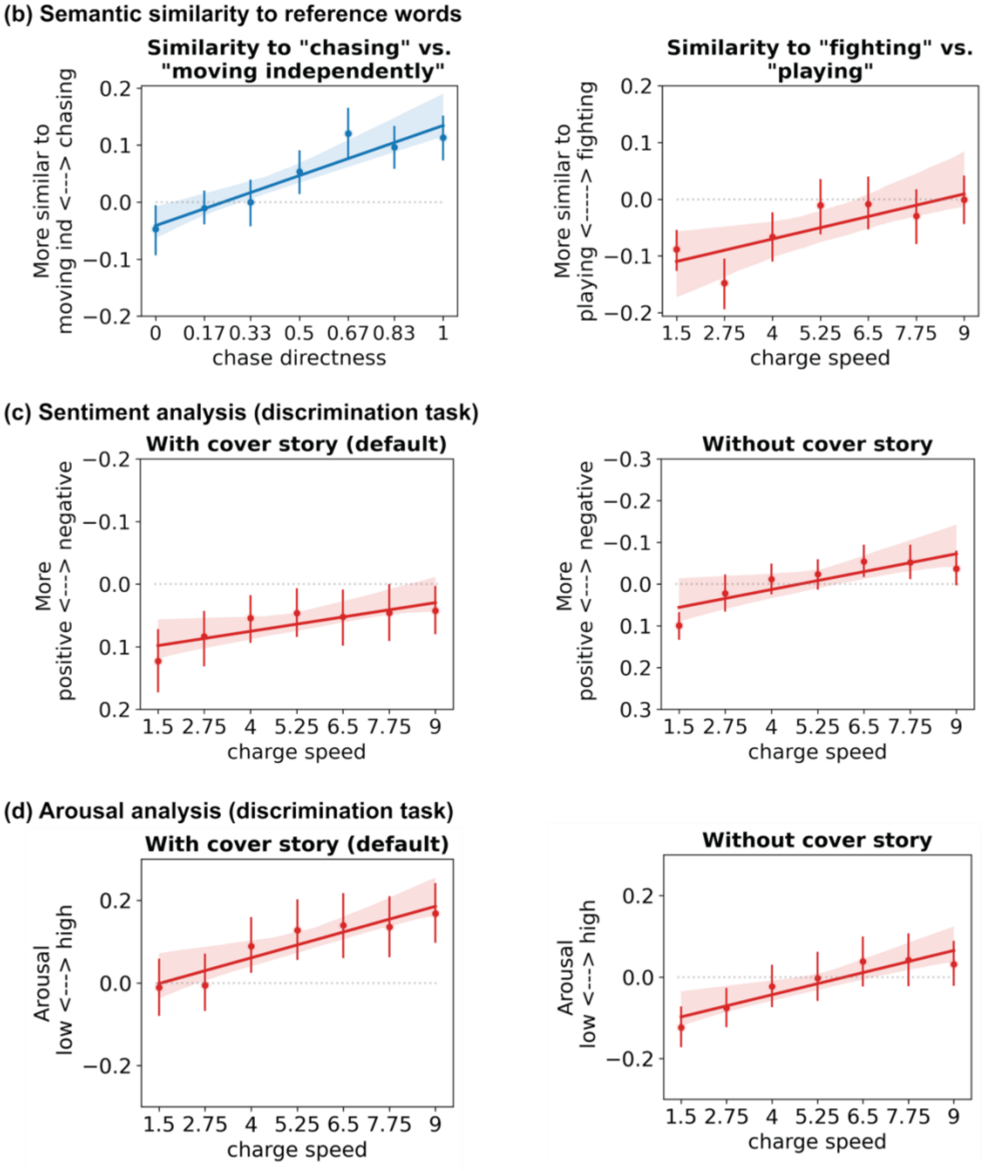

*Fig S1: Results from the free-text response pilot. (a) Unique verbs closest in semantic embedding space to the average of participants' descriptions of the detection (intended continuum: non-social ⟷ social) and discrimination (intended continuum: playing ⟷ fighting) animations, respectively. (b) Semantic similarity between average descriptions and intended anchor points. As expected, similarity to "chasing" relative to "moving independently" increases with chase directness for the detection stimuli (left), and similarity to "fighting" relative to "playing" increases with charge speed for the discrimination stimuli (right). (c) For the discrimination task, as charge speed increases, descriptions' negative sentiment increases relative to their positive sentiment - i.e., people perceived a trend in the expected positive ⟷ negative spectrum even if the exact words they used to describe the behaviors were not "playing" or "fighting". In the main version of the discrimination task presented with the cover story (left), we see an overall bias toward positive sentiment, potentially because of our "children in the playground" cover story, which likely induces a baseline expectation for positively valenced interactions. In an additional pilot without any cover story (right), this overall positive bias decreased (line shifts lower on the y-axis), while the pattern of increasing relative negative sentiment with increasing charge speed was maintained. (d) For the discrimination task, arousal too increased with charge speed, showing that the more negatively valenced responses also led to a higher arousal. Arousal was overall higher in the pilot batch with the cover story compared to the batch without the cover story. In spite of the similarity between valence and arousal in how they relate to charge speed, the effect of charge speed on arousal persisted even on including the arousal scores into the linear mixed effects models (see text).*

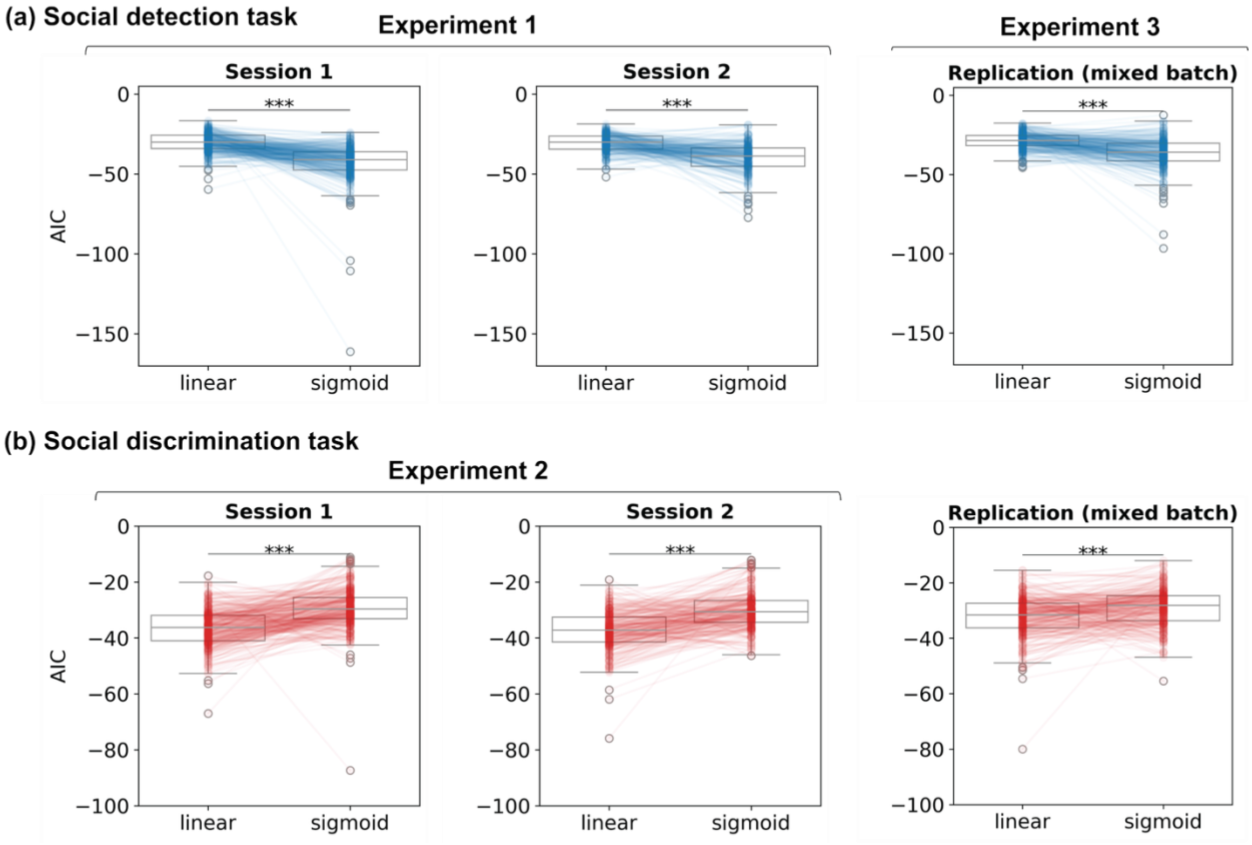


*Fig S2: Comparing linear vs. sigmoid fits for the (a) social detection and (b) social discrimination tasks, respectively using the Akaike Information Criterion (AIC). We see that for the social detection task, sigmoid fits are better as indicated by $AIC_{sigmoid} < AIC_{linear}$ (paired t-test: mean difference $AIC_{sigmoid} - AIC_{linear} = -12.24$, $-9.39$ and $-7.84$ left to right in panel (a)), whereas for the social discrimination task, linear fits are better as indicated by $AIC_{linear} < AIC_{sigmoid}$ (paired t-test: mean difference $AIC_{sigmoid} - AIC_{linear} = 7.21$, $7.3$ and $3.47$ left to right in panel (b)). *** = $p < .001$. Experiments 1, 2 and 3 refer to the detection, discrimination and mixed-task experiments, respectively.*

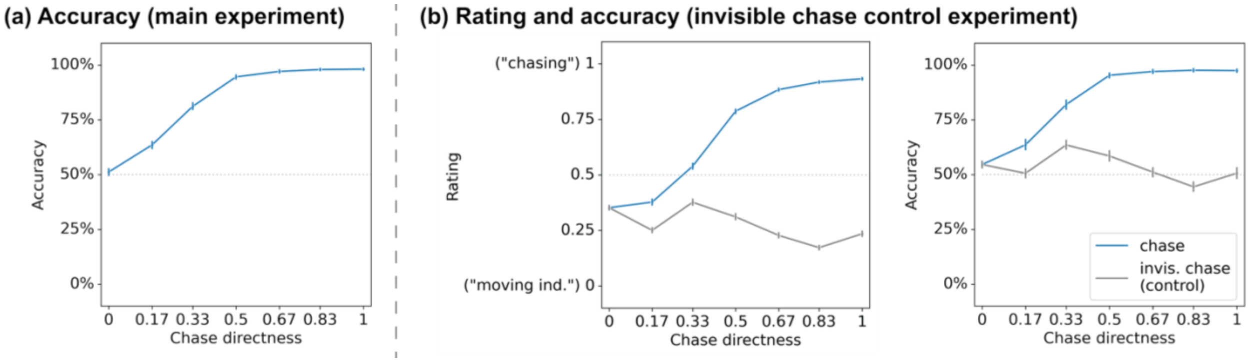


*Fig S3: (a) Predator identification accuracy increases as chases become more direct. Note that at directness 0 ("wander"), there is no contingent motion between the agents and therefore no correct answer; accuracy here reflects the percentage of trials on which participants identified the slower of the two agents as the predator. (True predators at all other levels of chase directness also moved slightly slower than their prey; therefore, if participants were using speed as a heuristic to identify predators, we would have seen above-chance "accuracy" in this condition, which results suggest was not the case). (b) Results from an auxiliary detection experiment with the additional control for correlated motion (with the invisible prey - see main text for details). Results show that chase directness increases socialness ratings and accuracy only in the true chase condition where both the predator and the prey are visible (blue line) and not in the invisible chase control (gray) condition. This shows that the ratings and accuracy do not simply represent sensitivity to correlated motion (which could have been a heuristic participants used to perform the chase detection task). N= (a) 312 and (b) 308. In the accuracy plots, the dotted line indicates chance performance. Error bars represent the 95% confidence interval.*

(a) Session 1, detection task

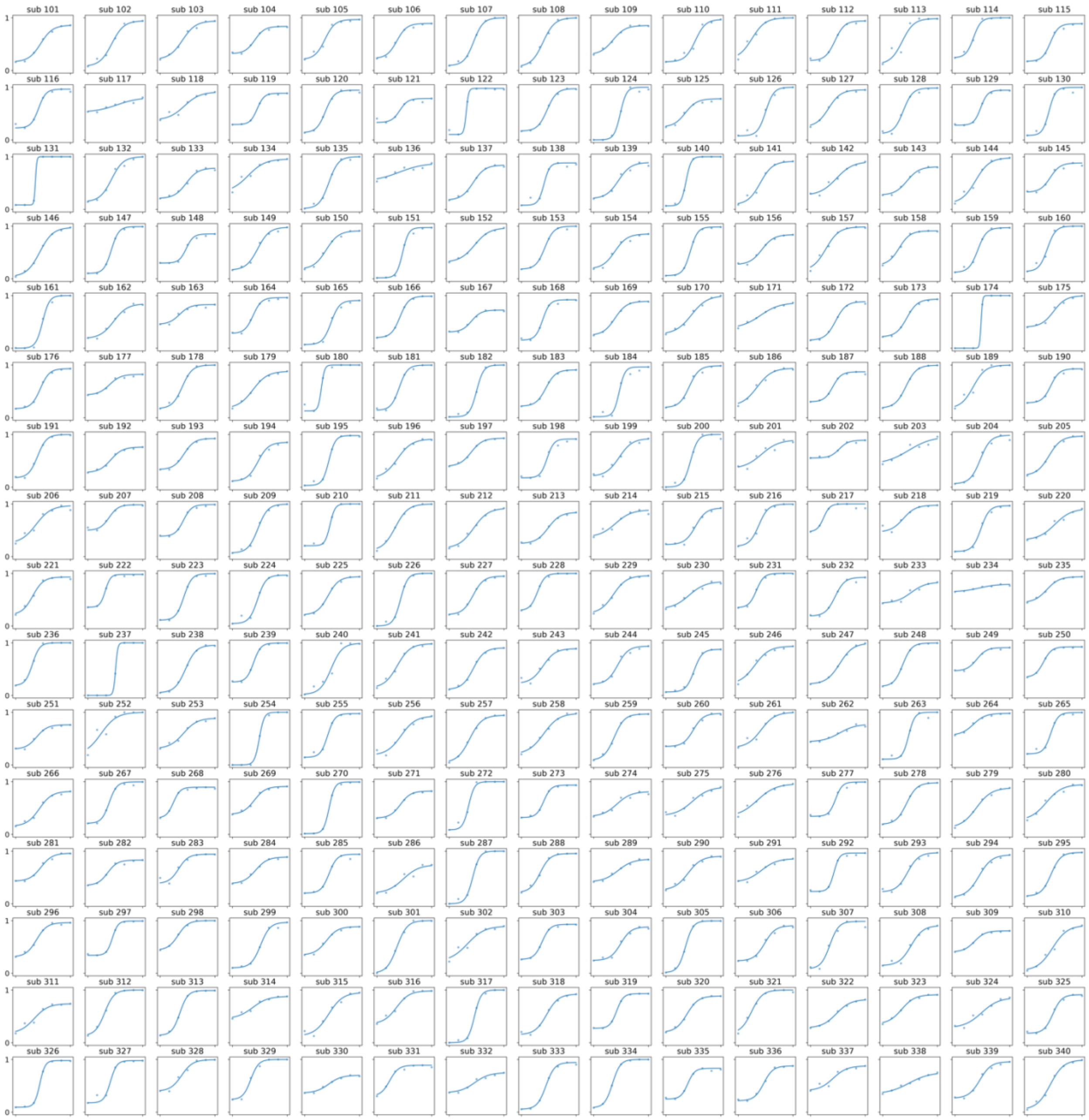

(b) Session 2, detection task

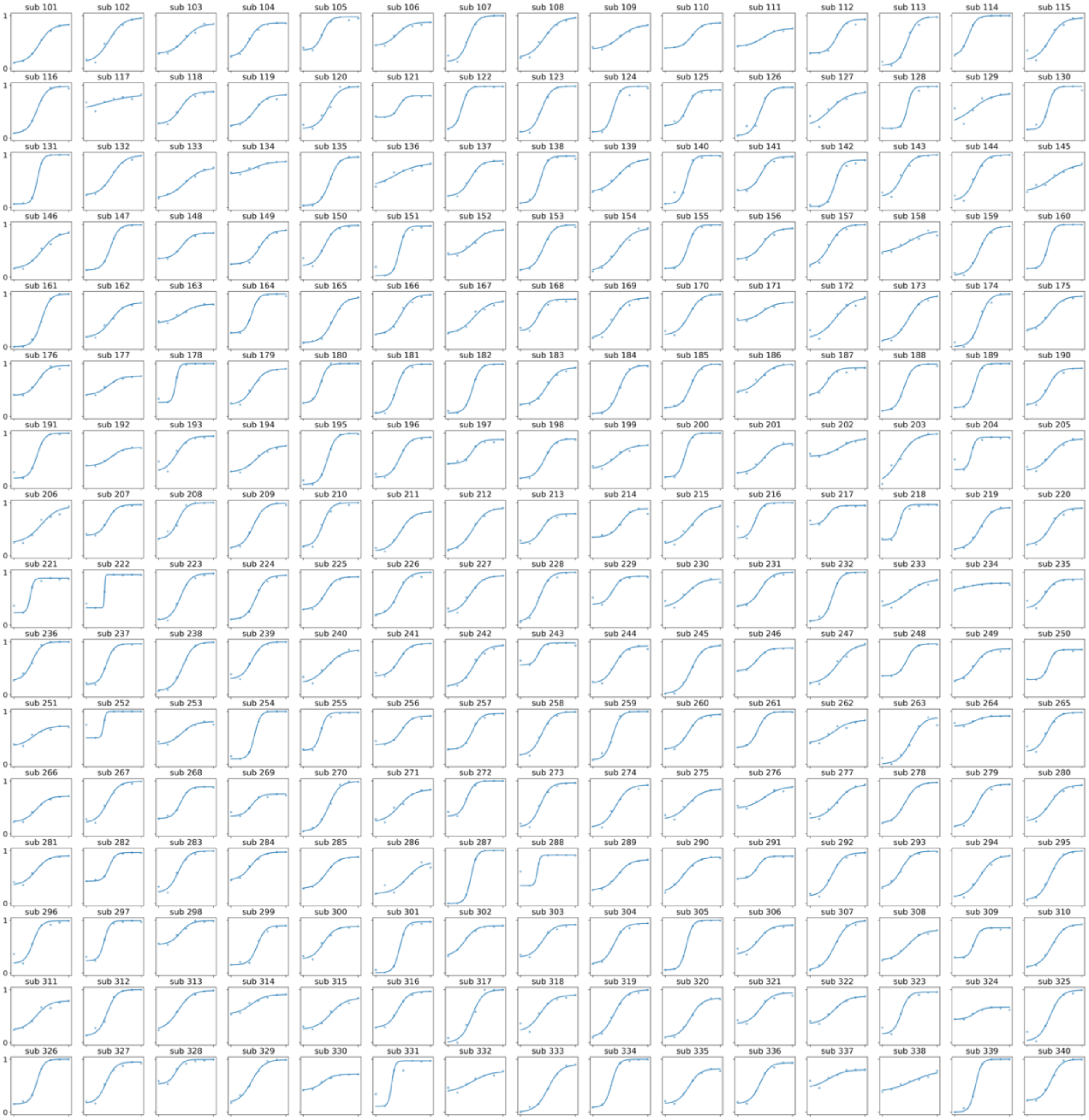


*Fig S4: Participant-level data from (a) session 1 and (b) session 2 of the detection experiment from all participants who participated in both sessions and rendered good quality data. Each sub-plot shows the mean ratings at each of the 7 chase directness levels (dots) and the best sigmoid fits (lines). x-axis = chase directness (objective motion attribute), y-axis = socialness rating.*

(a) Session 1, discrimination task

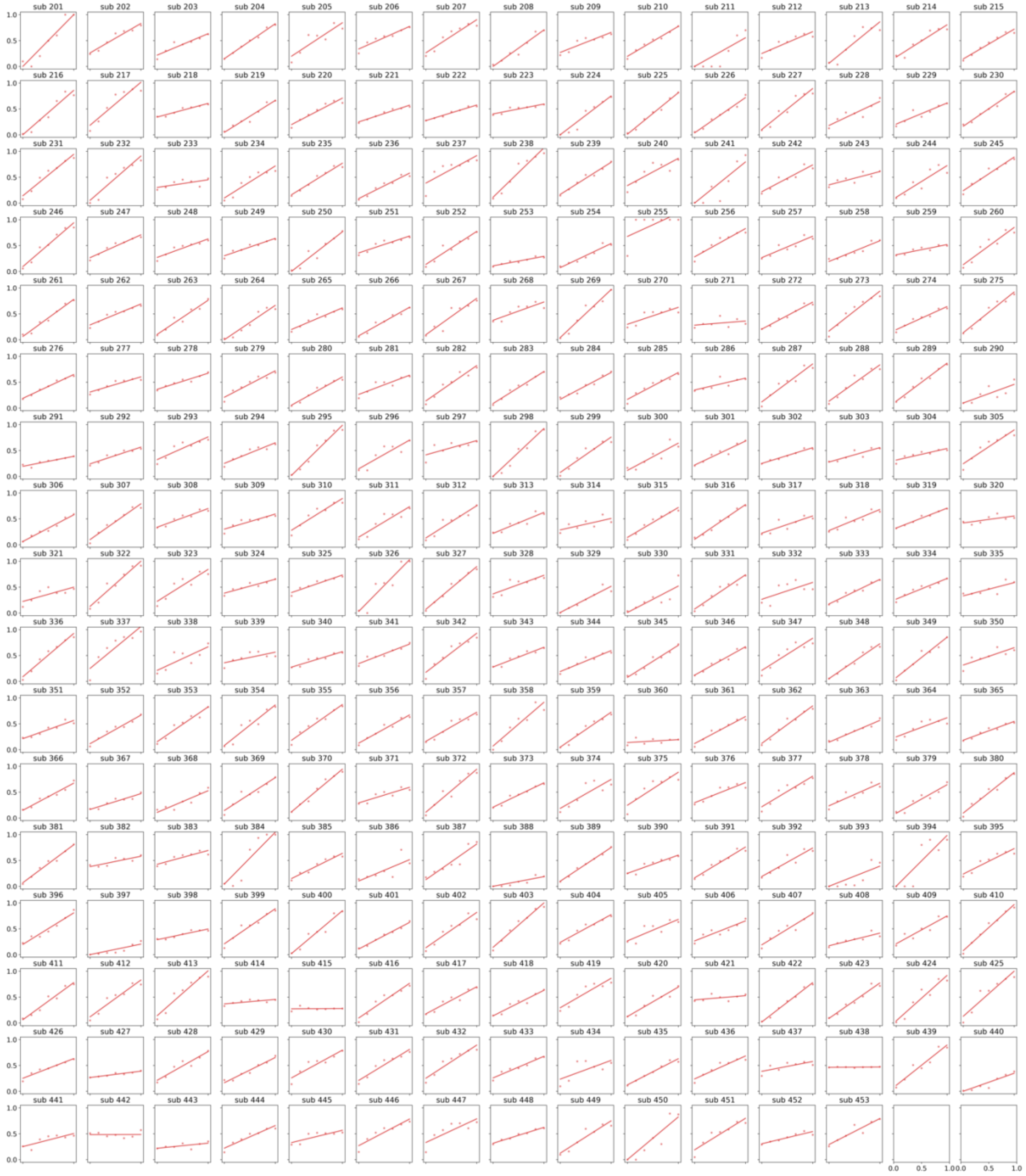

(b) Session 2, discrimination task

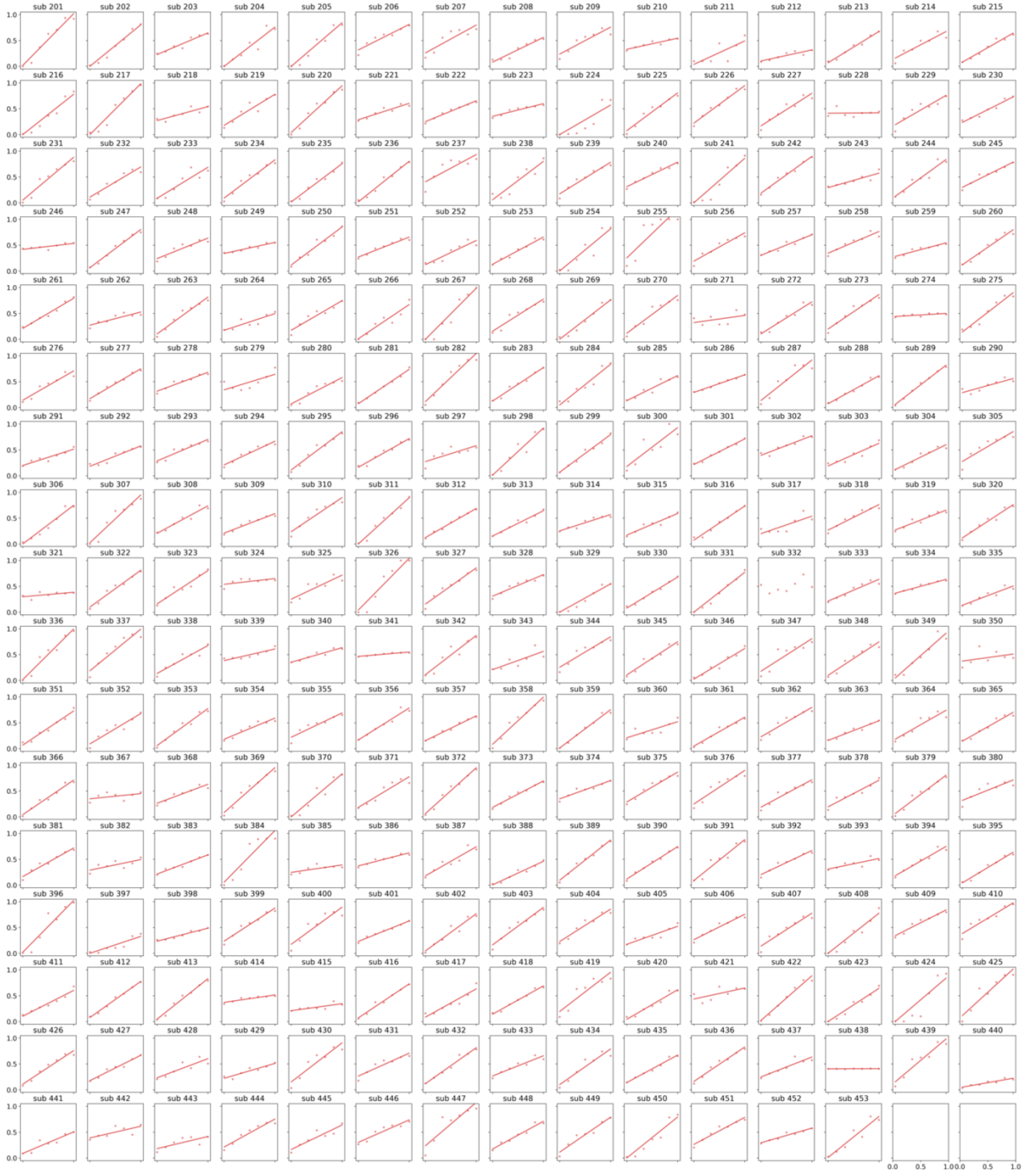


*Fig S5: Participant-level data from (a) session 1 and (b) session 2 of the discrimination experiment for all participants who participated in both sessions and rendered good quality data. Each sub-plot shows the*

*mean ratings (dots) and the best linear fits (lines). x-axis = charge speed (objective motion attribute), y-axis = subjective ratings of aggressiveness.*

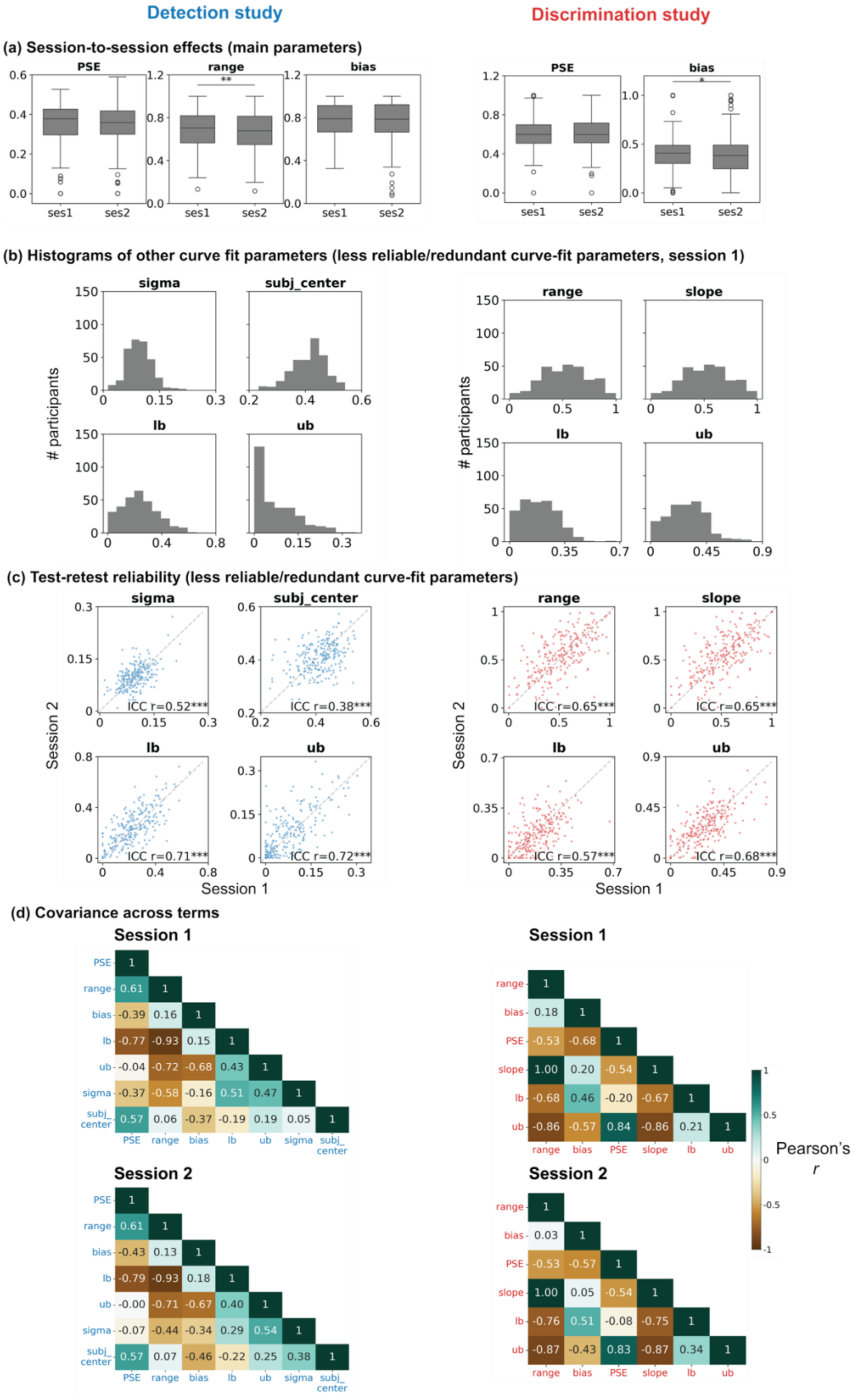

Detection study
Discrimination study
(a) Session-to-session effects (main parameters)
PSE
range
bias
**
*
ses1
ses2
(b) Histograms of other curve fit parameters (less reliable/redundant curve-fit parameters, session 1)
sigma
subj_center
lb
ub
range
slope
### participants
(c) Test-retest reliability (less reliable/redundant curve-fit parameters)
sigma
subj_center
lb
ub
range
slope
ICC r=0.52***
ICC r=0.38***
ICC r=0.71***
ICC r=0.72***
ICC r=0.65***
ICC r=0.65***
ICC r=0.57***
ICC r=0.68***
Session 2
Session 1
(d) Covariance across terms
Session 1
PSE 1
range 0.61 1
bias -0.39 0.16 1
lb -0.77 -0.93 0.15 1
ub -0.04 -0.72 -0.68 0.43 1
sigma -0.37 -0.58 -0.16 0.51 0.47 1
subj_center 0.57 0.06 -0.37 -0.19 0.19 0.05 1
PSE range bias lb ub sigma subj_center
Session 2
PSE 1
range 0.61 1
bias -0.43 0.13 1
lb -0.79 -0.93 0.18 1
ub -0.00 -0.71 -0.67 0.40 1
sigma -0.07 -0.44 -0.34 0.29 0.54 1
subj_center 0.57 0.07 -0.46 -0.22 0.25 0.38 1
PSE range bias lb ub sigma subj_center
Session 1
range 1
bias 0.18 1
PSE -0.53 -0.68 1
slope 1.00 0.20 -0.54 1
lb -0.68 0.46 -0.20 -0.67 1
ub -0.86 -0.57 0.84 -0.86 0.21 1
range bias PSE slope lb ub
Session 2
range 1
bias 0.03 1
PSE -0.53 -0.57 1
slope 1.00 0.05 -0.54 1
lb -0.76 0.51 -0.08 -0.75 1
ub -0.87 -0.43 0.83 -0.87 0.34 1
range bias PSE slope lb ub
Pearson's r

*Fig S6: Supplementary results for individual-difference analyses. (a) Curve parameters in session 1 vs. 2. Overall, the changes are small. In the detection task, range is slightly lower in session 2 (paired t-test: mean difference MD = 0.02, p = 0.008). In the discrimination task, in session 2, the bias towards aggressiveness is slightly lower (paired t-test: MD = –0.03, p = 0.03). (b) Histograms and (c) test-retest reliability of the curve parameters that showed lower test-retest reliability and/or were redundant with the main parameters of interest as indicated in the covariance matrix in b. (d) Pairwise correlations of all curve parameters across participants within each session. Each dot represents a participant. The covariance structure itself is highly stable across sessions. In both studies, the lower and upper biases (lb and ub, respectively) show a high retest reliability, but also correlate highly with range and PSE. For the sigmoid fit in the social detection task (left), sigma and center are somewhat independent of other terms, but their reliability is also lower (as indicated in a) than the 3 ultimately selected main parameters of interest. For the linear fit in the social discrimination task (right), slope shows good reliability but is redundant with range. *** = p < .001, * = p < .05.*

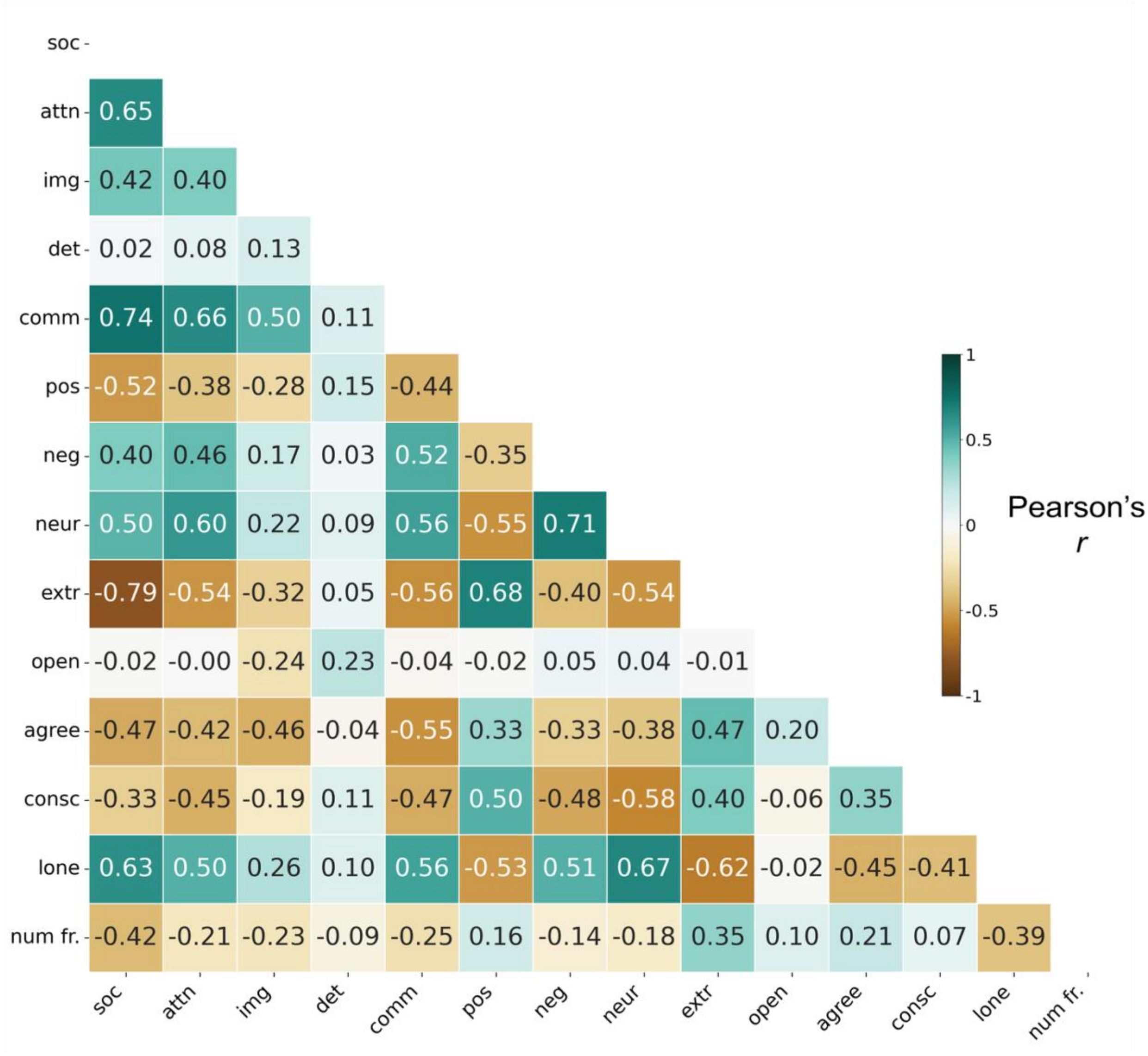


*Fig S7: Pearson correlation amongst the 14 trait dimensions. Abbreviations: Autism Quotient (AQ) questionnaire subscales – 'soc': social skill deficits, 'attn': attention-switching deficits, 'img': imagination deficits, 'det': heightened attention-to-detail and 'comm': communication deficits; Positive and Negative Affect Schedule (PANAS) subscales – 'pos': positive affect and 'neg': negative affect; Big 5 (NEO-FFI) subscales – 'neur': neuroticism, 'extr': extraversion, 'open': openness, 'agree': agreeableness and 'consc': conscientiousness; 'lone': loneliness score; '# fr.': number of friends.*

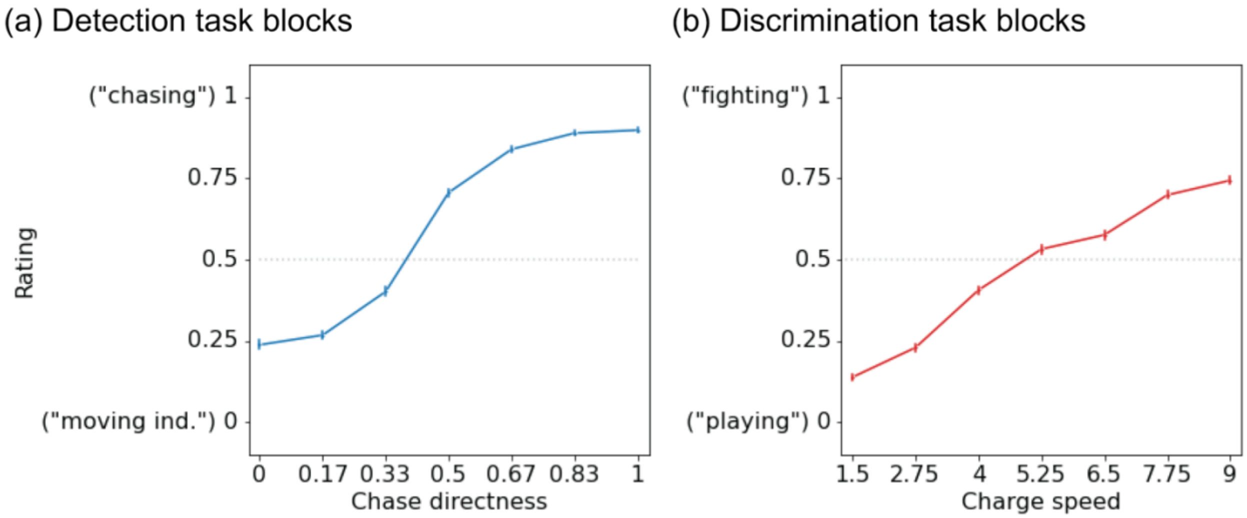


*Fig S8: Replication of the group-level results from the detection and discrimination task in an independent experiment (mixed-task experiment) where the same set of participants performed both tasks. (a) As the chases become more direct, the perceived socialness (y-axis) also increases. (b) As the speed at which agents charge at each other increases (x-axis), perception of an interaction becomes closer to "fighting" than "playing" (y-axis). N = 279. Error bars represent the 95% confidence interval.*
